# A subfunctionalization epistasis model to evaluate homeologous gene interactions in allopolyploid wheat

**DOI:** 10.1101/376731

**Authors:** Nicholas Santantonio, Jean-Luc Jannink, Mark E. Sorrells

## Abstract

Hybridization between related species results in the formation of an allopolyploid with multiple subgenomes. These subgenomes will each contain complete, yet evolutionarily divergent, sets of genes. Like a diploid hybrid, allopolyploids will have two versions, or homeoalleles, for every gene. Partial functional redundancy between homeologous genes should result in a deviation from additivity. These epistatic interactions between homeoalleles are analogous to dominance effects, but are fixed across subgenomes through self pollination. An allopolyploid can be viewed as an immortalized hybrid, with the opportunity to select and fix favorable homeoallelic interactions within inbred varieties. We present a subfunctionalization epistasis model to estimate the degree of functional redundancy between homeoallelic loci and a statistical framework to determine their importance within a population. We provide an example using the homeologous dwarfing genes of allohexaploid wheat, *Rht-1*, and search for genome-wide patterns indicative of homeoallelic subfunctionalization in a breeding population. Using the IWGSC RefSeq vl.0 sequence, 23,796 homeoallelic gene sets were identified and anchored to the nearest DNA marker to form 10,172 homeologous marker sets. Interaction predictors constructed from products of marker scores were used to fit the homeologous main and interaction effects, as well as estimate whole genome genetic values. Some traits displayed a pattern indicative of homeoallelic subfunctionalization, while other traits showed a less clear pattern or were not affected. Using genomic prediction accuracy to evaluate importance of marker interactions, we show that homeologous interactions explain a portion of the non-additive genetic signal, but are less important than other epistatic interactions.

## 2 Introduction

Whole genome duplication events are ubiquitous in the plant kingdom. The impact of these duplications on angiosperm evolution was not truly appreciated until the ability to sequence entire genomes elucidated their omnipresence (Soltis *et al*., 2009). Haldane (1933), postulated that single gene duplication allowed one copy to diverge through mutation while metabolic function was maintained by the other copy. Ohno (1970) reintroduced this hypothesis, and it has since been validated both theoretically (Ohta, 1987; Walsh, 1995; Lynch and Conery, 2000), and empirically (Blanc and Wolfe, 2004; Duarte *et al*., 2005; Liu *et al*., 2011; Assis and Bachtrog, 2013). The duplicated gene hypothesis does not, however, generally explain the apparent advantage of duplicating an entire suite of genes. The necessity of genetic diversity for plant populations to survive and adapt to divergent or changing environments may help to explain this pervasive phenomenon.

The need for gene diversity can become more immediate in plants than in animals, where the latter can simply migrate to “greener pastures” when conditions become unfavorable. Plants lack substantial within generation mobility and must therefore change gene expression to cope with changing environmental conditions. Many species maintain gene diversity through alternate splicing, but this has been shown to be less common in plants than in other eukaryotes (Nagasaki *et al*., 2005). Whole genome duplication can generate the raw materials for the maintenance of genetic diversity (Wendel, 2000; Adams and Wendel, 2005). Gault *et al.* (2018) demonstrated that similar sets of duplicated genes were preserved in two related genera, *Zea* and *Tripsacum*, millions of years after a shared paleopolyploidization event. This conserved pattern in purifying selection suggests that, at least for some genes, there is a clear advantage to maintaining two copies.

The union of two complete, yet divergent, genomes during the formation of an allopolyploid introduces manifold novel gene pathways that can specialize to specific tissues or environments (Blanc and Wolfe, 2004). Similar to diploid hybrids, the formation of an allopolyploid results in a homogeneous population, but heterozygosity is maintained across homeologous sites rather than homologous sites. Unlike diploid hybrids that lose heterozygosity in subsequent generations, the homeoallelic heterozygosity is fixed through selfing in the allopolyploid. Mac Key (1970) postulated a trade off between new-creating (allogamous) and self preserving (autogamous) mating systems, where allopolyploids favor self pollination to preserve diverse sets of alleles across their subgenomes. As such, an allopolyploid may be thought of as an immortalized hybrid, with heterosis fixed across subgenomes (Ellstrand and Schierenbeck, 2000; Feldman *et al*., 2012). While still hotly debated, evidence is mounting that allopolyploids exhibit a true heterotic response as traditional hybrids have demonstrated (Wendel, 2000; Adams and Wendel, 2005; Chen, 2010, 2013).

Birchler *et al*., (2010) note that newly synthesized allopolyploids often outperform their sub-genome progenitors, and that the heterotic response appears to be exaggerated in wider inter-specific crosses. This seems to hold true even within species, where autopolyploids tend to exhibit higher vigor from wider crosses (Bingham *et al*., 1994; Segovia-Lerma *et al*., 2004). The overwhelming prevalence of allopolyploidy to autopolyploidy in plant species (Soltis and Soltis, 2009) may suggest that it is the increase in allelic diversity *per se* that is the primary driver for this observed tendency toward genome duplication. Instead of allowing genes to change function after a duplication event, alleles may develop novel function prior to their reunion during an allopolyploidization event. The branched gene networks of the allopoloyploid may provide the organism with the versatility to thrive in a broader ecological landscape than those of its sub-genome ancestors (Mac Key, 1970; Ellstrand and Schierenbeck, 2000; Osborn *et al*., 2003).

Subfunctionalization and neofunctionalization are often described as distinct evolutionary processes. Neofunctionalization implies the duplicated genes have completely novel, non-redundant function (Ohno, 1970). Subfunctionalization is described as a partitioning of ancestral function through degenerative mutations in both copies, such that both genes must be expressed for physiological function (Stoltzfus, 1999; Force *et al*., 1999; Lynch and Force, 2000). However, barring total functional gene loss, many mutations will have some quantitative effect on protein kinetics or expression (Zeng and Cockerham, 1993). Duplicated genes will demonstrate some quantitative degree of functional redundancy until the ultimate fate of neofunctionalization (i.e. complete additivity) or gene loss (pseudogenization) of one copy. It has been proposed that essentially all neofunctionalization processes undergo a subfunctionalization transition state (Rastogi and Liberles, 2005).

If the mutations occur before the duplication event, as in allopolyploidy, the two variants are unlikely to have degenerative mutations. Instead, they may have differing optimal conditions in which they function or are expressed. The advantage of different variants at a single locus (alleles; Allard and Bradshaw, 1964) or at duplicated loci (homeoalleles; Mac Key, 1970) can result in greater plasticity to environmental changes. Allopolyploidization has been suggested as an evolutionary strategy to obtain the genic diversity necessary for invasive plant species to adapt to the new environments they invade (Ellstrand and Schierenbeck, 2000; te Beest *et al*., 2011).

Adams *et al*., (2003) showed that some homeoallelic genes in cotton were expressed in an organ specific manner, such that expression of one homeolog effectively suppressed the expression of the other in some tissues. These results have since been confirmed in other crops such as wheat (Pumphrey *et al*., 2009; Akhunova *et al*., 2010; Feldman *et al*., 2012; Pfeifer *et al*., 2014), and evidence for neofunctionalization of homeoallelic genes has been observed (Chaudhary *et al*., 2009). Differential expression of homeologous gene transcripts has also been shown to shift upon challenge with heat, drought (Liu *et al*., 2015) and salt stress (Zhang *et al*., 2016) in wheat, as well as water submersion and cold in cotton (Liu and Adams, 2007).

Common wheat (*Triticum aestivum*) provides an example of an allopolyploid that has surpassed its diploid ancestors in its value to humans as a staple source of calories. Hexaploid wheat has undergone two allopolyploid events, the most recent of which occurred between 10 and 400 thousand years ago, adding the D genome to the A and B genomes (Marcussen *et al*., 2014). The gene diversity provided by these three genome ancestors may explain why allohexaploid wheat has adapted from its source in southwest Asia to wide spread cultivation around the globe (Dubcovsky and Dvořák, 2007; Feldman and Levy, 2012).

In the absence of outcrossing in inbred populations, selection can only act on individuals, changing their frequency within the population. If the selection pressure changes (e.g. for modern agriculture), combinations of homeoalleles within existing individuals may not be ideal for the new set of environments and traits. This presents an opportunity for plant breeders to capitalize on this feature of allopolyploids by making crosses to form new individuals with complementary sets of homeoalleles. Many of these advantageous combinations have likely been indirectly selected throughout the history of wheat domestication and modern breeding.

Dominance of homeologous genes is known to exist in wheat. For example, a single dominant red allele at any of the three homeologous kernel color genes on 3A, 3B, and 3D will confer a red kernel color (Allan and Vogel, 1965; Metzger and Silbaugh, 1970). Another crucial example involves the two homeologous dwarfing genes (Allan *et al*., 1959; Gale *et al*., 1975; Gale and Marshall, 1976; McVittie *et al*., 1978) important in the Green Revolution, which implemented semi-dwarf varieties to combat crop loss due to nitrogen application and subsequent lodging. These genes have been shown to exhibit a quantitative semi-dominant reponse (Börner *et al*., 1996). We discuss this example in detail, and use it as a starting point to justify the search for quantitative homeologous interactions genome-wide. While the effect of allopolyploidy has been demonstrated at both the transcript level and whole plant level, we are unaware of attempts to use genome-wide homeologous interaction predictors to model whole plant level phenotypes such as growth, phenology and grain yield traits.

Using a soft winter wheat breeding population, we demonstrate that epistatic interactions account for a significant portion of genetic variance and are abundant throughout the genome. Some of these interactions occur between homeoallelic regions and we demonstrate their potential as targets for selection. If advantageous homeoallelic interactions can be identified, they could be directly selected to increase homeoallelic diversity, with the potential to expand the environmental landscape to which a variety is adapted. We hypothesize that the presence of two evolutionarily divergent genes with partially redundant function leads to a less-than-additive gene interaction, and introduce this as a subfunctionalization model of epistasis.

## 3 Subfunctionalization Epistasis

We generalize the duplicate factor model of epistasis from Hill *et al*., (2008), by introducing a subfunctionalization coefficient *s*, that allows the interaction to shift between the duplicate factor and additive models. Let us consider an ancestral allele with an effect *a.* Through mutation, the effect of this locus is allowed to diverge from the ancestral allele to have effects *a*\* and *ã* in the two descendant species. When the two divergent loci are brought back together in the same nucleus, the effect of combining these becomes *s*(*a*\* + *ã*) (Figure 1).

**Figure 1:**
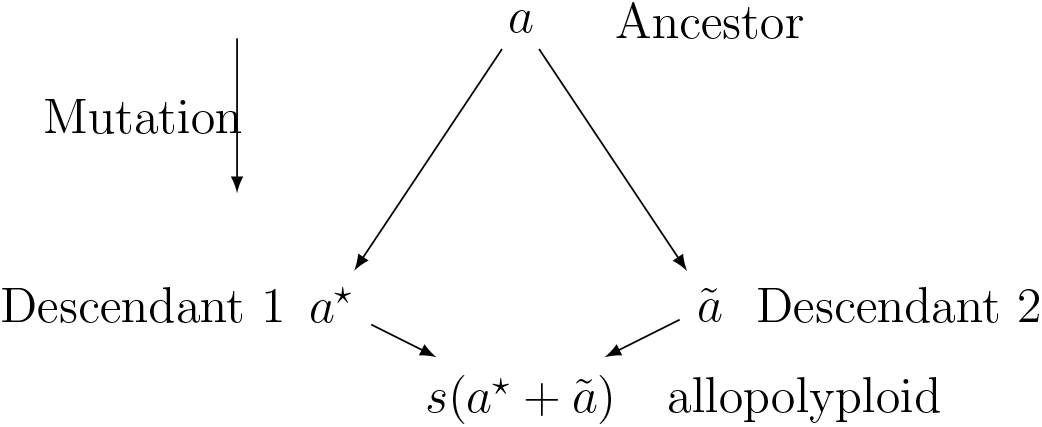
Diagram of subfunctionalization where *a* is the effect of a functional allele, *a*\* and *ã* are the effects of the descendant alleles, and *s* is the subfunctionalization coefficient.

Values of *s* < 1, indicate a less-than additive epistasis (Eshed and Zamir, 1996), in this case, resulting from redundant gene function. When *s* = 1/2, and *a*\* = *ã*, the descendant alleles have maintained the same function and the duplicate factor model is obtained. As *s* exceeds 1/2, the descendant alleles diverge in function (i.e. subfunctionalization), until *s* reaches 1, implying that the two genes evolved completely non-redundant function (i.e neofunctionalization). At the point where *s* = 1, the effect becomes completely additive.

For values of *s* > 1/2, the benefit of multiple alleles is realized in a model analogous to overdominance in traditional hybrids. As alleles diverge they can pick up advantageous function under certain environmental conditions. The homeo-heterozygote then gains an advantage if it experiences conditions of both adapted homeoalleles. Values of *s* < 1/2 may indicate allelic interference (Herskowitz, 1987), or genomic shock (McClintock, 1984), a phenomenon that has been observed in many newly formed allopolyploids (Comai *et al*., 2003). Allelic interference, also referred to as dominant negative mutation, can result from the formation of non-functional homeodimers, while homodimers from the same ancestor continue to function properly. This interference effectively reduces the number of active dimers by half (Herskowitz, 1987; Veitia, 2007).

### 3.1 Epistasis models

Let us consider the two locus model, with loci *B* and *C.* Using the notation of Hill *et al*., (2008), the expected phenotype, *E*[*y*], is modeled as

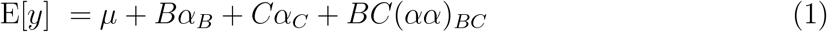

where *B* and *C* are the marker allele scores, BC is the pairwise product of those scores, *α_B_* and *α_C_* are the additive effects of the B and C loci and (*αα*)_*BC*_ is the interaction effect.

We revisit two epistatic models, the “Additive × Additive Model without Dominance or Interactions Including Dominance” (called “Additive × Additive” hence forth) and the “Duplicate Factor” considered by Hill, Goddard and Visscher (Hill *et al*., 2008) that are relevant for this discussion. Omitting the heterozygous classes and letting *a* be the effect on the phenotype,

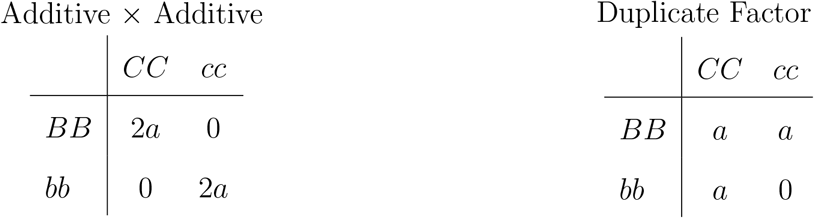

We propose a generalized Duplicate Factor epistatic model to estimate the degree of gene functional redundancy, or subfunctionalization.

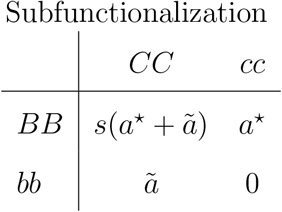

When markers are coded {0, 1} for presence of the functional allele, the deviation from the additive expectation, *δ*, is estimated by (*αα)_BC_*. *δ* can then be used to calculate the subfunctionalization coefficient, 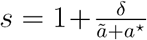 (Figure 2). The least squares expectation of additive and epistatic effects is then

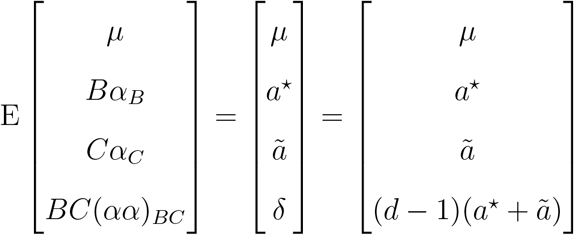

**Figure 2:**
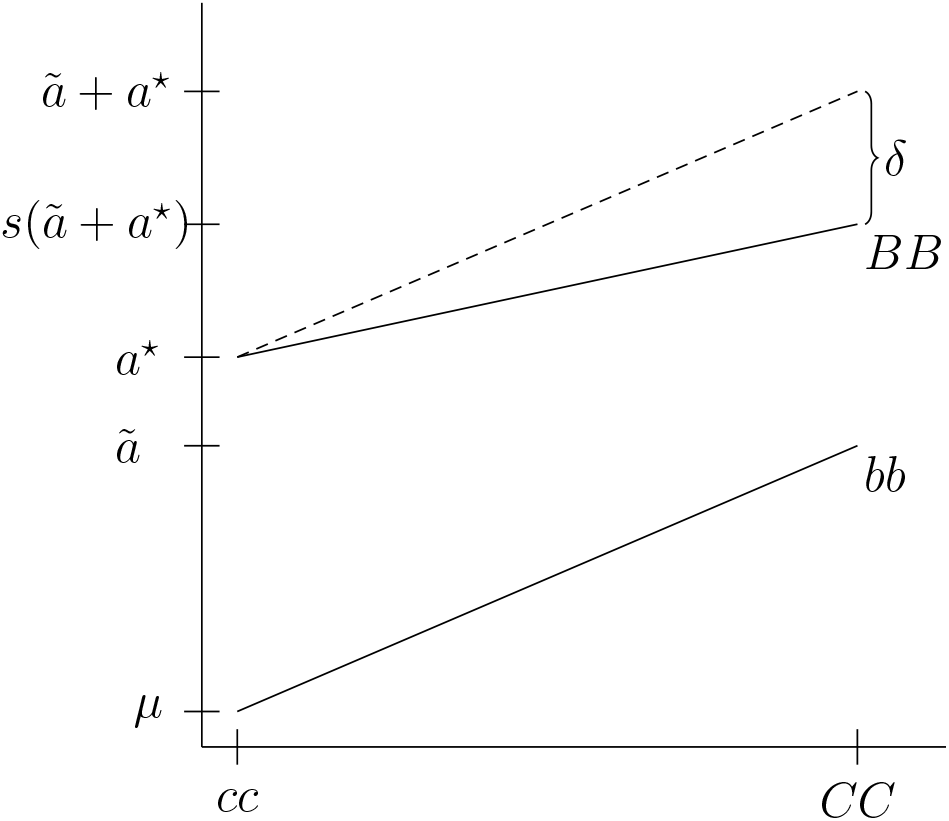
Epistatic interaction of two loci, *B* and *C*, with the expected effects for the {0, 1} parameterization. <5 indicates the deviation of the *BBCC* genotype from an additive model for the {0, 1} parameterization, where 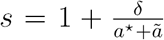 The dotted line indicates the expectation under the additive model.

### 3.2 Epistatic contrasts

Epistatic interaction predictors must be formed from marker scores in order to estimate interaction parameters. These interaction predictors are typically calculated as the pairwise product of the genotype scores for their respective loci. This can lead to ambiguity in the meaning of those interaction effects depending on how the marker scores are coded. Different marker parameterizations can center the problem at different reference points (i.e. different intercepts), and can scale the predictors based on allele or genotype effects (i.e. different slopes).

When loci *B* and *C* are coded as {−1, 1} for inbred genotypes, including the product of the marker scores, *BC*, corresponds to the Additive × Additive model (Table 1). Changing the reference allele at either locus does not change the magnitude of effect estimates but will change their signs. Using {0, 1} coding, *BC* corresponds to the subfunctionalization model and estimates *δ* directly. For this coding scheme, the magnitude and sign can change depending on the reference allele at the two loci. This highlights one of the difficulties of effect interpretation, as it is not clear which marker orientations should be paired. That is, which allele should be *B* as opposed to *b*, and which should be *C* as opposed to *c*? Marker alleles can be oriented to have either all positive or all negative additive effects, but the question remains: which direction should the more biologically active allele have on the phenotype?

**Table 1:** Epistatic interaction score tables resulting from the products of marker scores using {−1, 1} and {0, 1} parameterizations for inbreds.

| | $CC$ | $cc$ | | $CC$ | $cc$ |
| --- | --- | --- | --- | --- | --- |
| $BB$ | 1 | -1 | $BB$ | 1 | 0 |
| $bb$ | -1 | 1 | $bb$ | 0 | 0 |

Marker scores are typically assigned as either presence (or absence) of the reference, major, or minor allele, which may or may not be biologically relevant. While it has been noted that the two different marker encoding methods do not result in the same contrasts of genotypic classes (He *et al*., 2015; Martini *et al*., 2016, 2017), coding does not affect the least squares model fit (Zeng *et al*., 2005; Álvarez-Castro and Carlborg, 2007). Álvarez-Castro and Carlborg (2007) show that there exists a linear transformation to shift between multiple parameterizations using a change-of-reference operation (see Appendix 3). This is convenient because all marker orientation combinations can be easily generated by changing the effect signs of a single marker orientation fit for the {−1, 1} marker coding. These effect estimates can subsequently be transformed to the {0, 1} coding effect estimates using the change-of-reference operation for all marker orientation combinations.

This transformation does not hold when marker effects are considered random, where the interaction effect is subject to differential shrinkage depending on the marker coding and orientation (Martini *et al*., 2017, 2018). As such, orienting markers to capture functional allele relationships may be crucial for optimizing genomic prediction including epistasis. We make an attempt to orient markers based solely on estimated fixed marker additive effects, with the assumption that that homeoalleles with similar additive effects are functionally similar. Other attempts at marker orientation have included orienting markers to maximize the interaction effect magnitude and including interaction predictors from all possible marker orientations (Martini *et al*., 2017). The former is biased toward selecting interaction predictors with a high joint frequency, whereas the latter suffers from a high degree of linear dependency.

## 4 Materials and Methods

### 4.1 RIL population

A bi-parental recombinant inbred line (RIL) population of 158 lines segregating for two dwarfing genes was used to illustrate an epistatic interaction between the well known homeologous genes on chromosomes 4B and 4D, *Rht-Bl* and *Rht-Dl*, important in the Green Revolution (Allan *et al*., 1959; Gale *et al*., 1975; Gale and Marshall, 1976; McVittie *et al*., 1978). Two genotyping by sequencing (GBS) markers linked to these genes were used to track the segregating mutant (*b* and *d*) and wildtype (*B* and *D*) alleles. Only one test for epistasis between these two markers was run. This homeologous marker pair was denoted RIL_Rhtl. Details of the population can be found in Appendix 1.

### 4.2 CNLM population

The Cornell small grains soft winter wheat breeding population (CNLM) was used to investigate the importance of homeologous gene interactions in a large adapted breeding population. The dataset and a detailed description of the CNLM population can be found in Santantonio *et al*., (2018b). Briefly, the dataset consists of 1,447 lines evaluated in 26 environments around Ithaca, NY. Because the data were collected from a breeding population, only 21% of the genotype/environment combinations were observed, totaling 8,692 phenotypic records. Standardized phenotypes of four traits, grain yield (GY), plant height (PH), heading date (HD) and test weight (TW) were recorded. All lines were genotyped with 11,604 GBS markers aligned to the International Wheat Genome Sequencing Consortium (IWGSC) RefSeq vl.0 wheat genome sequence of ‘Chinese Spring’ (IWGSC, 2018, accepted), and subsequently imputed.

### 4.3 Homeologous marker sets

Using the IWGSC RefSeq vl.0 ‘Chinese Spring’ wheat genome sequence (IWGSC, 2018, accepted), homeologous sets of genes were constructed by aligning the annotated coding sequences (vl.0) back onto themselves. The known 4A, 5A, and 7B translocation in wheat (Devos *et al*., 1995) was ignored for simplicity in this study, but could easily be accounted for by allowing homeologous pairs across these regions. The resulting 23,796 homeologous gene sets, comprised of 18,184 triplicate and 5,612 duplicate gene sets, sampled roughly 59% of the gene space of hexaploid wheat. Additional details on homeologous gene alignment can be found in Appendix 2. Each homeologous gene was then anchored to the nearest marker by physical distance (Supplementary Figures S1 and S2), and homeologous sets of markers were constructed from the anchor markers to each homeologous gene set. Redundant marker sets due to homeologous genes anchored by the same markers were removed, resulting in 6,142 triplicate and 3,985 duplicate marker sets for a total of 10,127 unique homeologous marker sets. These marker sets (denoted ‘Homeo’) were then used to calculate marker interaction scores as pairwise products of the marker score vectors.

As a control, two additional marker sets were produced by sampling the same number of duplicate and triplicate marker sets as the Homeo set. These markers sets were sampled either from chromosomes within a subgenome (Within, e.g. markers on 1A, 2A and 3A), or across non-syntenic chromosomes of different subgenomes (Across, e.g. markers on 1A, 2B and 3D). Samples were taken to reflect the same marker distribution of the Homeo set with regard to their native genome, which has a larger proportion of D genome markers relative to their abundance. Note that three-way homeologous interactions have equal proportions of markers belonging to the A, B and D genomes, whereas D genome markers only account for 13% of all markers in the CNLM population (Santantonio *et al*., 2018b).

### 4.4 Determining marker orientation

For each homeologous marker set, additive homeologous marker effects and their multiplicative interaction effects were estimated as fixed effects in the following linear mixed model while correcting for background additive and epistatic effects. The {–1, 1} marker parameterization was used for fixed marker additive and interaction effect estimates.

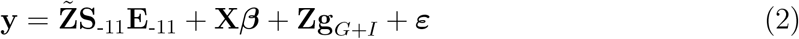

where **X** is the design matrix, *β* is the vector of fixed environmental effects, and **Z** is the line incidence matrix. **S**_-11_ is the matrix of genotype marker scores and interactions for each genotype class, while **E**_-11_ is the fixed additive and interaction effects that need estimated (Appendix 3). 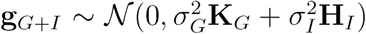 is the incidence matrix for the two-or three-way genotype of each homeologous marker set. **Z** and 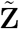 differ in that the former links observations to a specific line, whereas the latter links observations to one of the two-or three-way genotype classes for the homeologous marker set. The background genetic effects were assumed to be 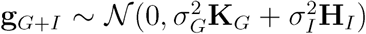 with population parameters previously determined (Zhang *et al*., 2010). The additive and epistatic covariances, **K**_*G*_ and **H**_*I*_, were calculated as described in VanRaden (2008, method I) and Martini *et al*., (2016, equation 9), respectively. This weighted covariance matrix was used to reduce computational burden associated with estimating two variance components in the same fit.

A Wald test was used to obtain a p-value for marker additive and interaction effects. All marker orientation combinations were generated by changing the estimated effect signs, and then transformed to the {0, 1} marker effect estimates using the change-of-reference operation (Álvarez-Castro and Carlborg, 2007). Only marker orientations with all positive or all negative additive effects were considered. It should be noted that the marker orientation has no effect on the p-value, as they are linear transformations of one another.

Markers were oriented to have minimized the difference (or variance for three-way sets) of the additive main effects while maximizing the mean of the absolute values of the additive main effects. This orientation, which we denote ‘low additive variance high additive effect’ (LAVHAE), assumes that marker alleles with similar effects are functionally similar. Only additive effects were used to select the marker orientation to keep from systematically selecting marker orientations with a specific interaction pattern. Three other marker orientation schemes were also investigated by orienting markers to either have all positive (POS) effects, all negative (NEG) effects, or to maximize the variance of the additive and interactions effects (‘high total effect variance’, HTEV).

### 4.5 Additive only simulated controls

Marker effect and interaction estimates using either {0, 1} or {−1, 1} marker parameterizations are not orthogonal, so care must be taken when interpreting the direction and magnitude of the effects estimates. The positive covariance between the marker scores and their interaction leads to a multicollinearity problem, and results in a negative relationship between additive and interaction effects if both additive effects are oriented in the same direction. To determine if the negative relationship between the additive and epistatic effects was greater than expected due to multicollinearity, a new phenotype with no epistatic effects was simulated from the data for each trait. The estimate of the marker variance was calculated from the additive genetic variance estimate as 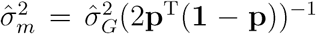, where **p** is the vector of marker allele frequencies. Then for each trait, a new additive phenotype was simulated as 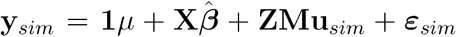 where the trial effect estimates from Santantonio *et al*., (2018b, equation 2) were used for 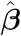, **M** is the matrix of marker scores, **u**_*sim*_ was sampled from 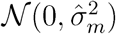 and *ε* was sampled from 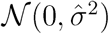. A Kolmogorov-Smirnov (KS) test was used to determine if the distribution of the estimated interaction effects from the actual data differed from the distribution of effects estimated from simulated data. An additional simulated phenotype was also produced by first permuting each column of M to remove any effects due to linkage disequilibrium (LD) structure.

### 4.6 Genomic prediction

To determine the importance of epistatic interactions to the predictability of a genotype, a genomic prediction model was fit as

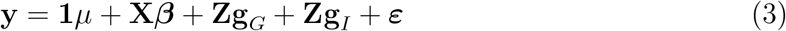

where **1**_*n*_ is a vector of ones, *μ* is the general mean. The random vectors of additive genotype, epistatic interactions, and errors were assumed to be distributed as 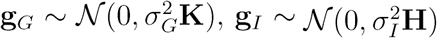 and 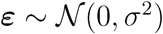, respectively.

The additive covariance matrix, **K,** was calculated using VanRaden (2008), method **I.** The epistatic covariance matrix **H** was calculated either as defined by Jiang and Reif (2015, equation 5) and Martini *et al*., (2016, equation 9) to model all pairwise epistatic interactions using {–1, 1} coding (Pairwise), or in a similar fashion as **K** for oriented marker sets, where only unique products of marker variables were included instead of the marker variables. For the latter, the matrix was scaled with the sum of the joint marker variances as (2**q**^τ^(1 –**q**))^−1^, where **q** is the joint frequency of individuals containing both the non-reference marker alleles. Three-way marker products were included if they were unique from the additive and pairwise product predictors.

A small coefficient of 0.01, was added to the diagonals of the covariance matrix to recover full rank lost in centering the matrix of scores prior to calculating the covariance. Five-fold cross validation was performed by randomly assigning individuals to one of five folds for 10 replications. Four folds were used to train the model and predict the fifth fold for all five combinations. All models were fit to the same sampled folds so that models would be directly comparable to one another and not subject to sampling differences. Prediction accuracy was assessed by collecting genetic predictions for all five folds, then calculating the Pearson correlation coefficient between the predicted genetic values for all individuals and a “true” genetic value. The “true” genetic values were obtained by fitting a mixed model to all the data with fixed effects for environments and a random effect for genotypes, assuming genotype independence with a genetic covariance **I.**

Increase in genomic prediction accuracy from the additive model was used as a proxy to assess the relative importance of oriented marker interaction sets. To determine the proportion of non-additive genetic signal attributable to each interaction set, the ratio of the prediction accuracy increase from the additive model using the interaction set (Homeo, Within and Across) to the prediction accuracy increase from the additive model modeling all pairwise epistatic interactions (Pairwise) was used for comparison of models. The percentage of non-additive predictability was calculated as follows for each interaction set.

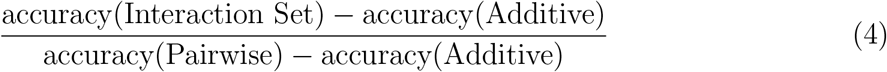

### 4.7 Software

ASReml-R (Gilmour, 1997; Butler, 2009) was used to fit all mixed models. BLAST+ (Camacho *et al*., 2009) was used for coding sequence alignment. All additional computation, analyses and figures were made using base R (R Core Team, 2015) implemented in the Microsoft Open R environment 3.3.2 (Microsoft, 2017) unless noted otherwise. Figures 1 and 2 were created using the ‘tikz’ package (Tantau, 2018) for F1¾X. Figure 4 was made with the ‘circlize’ R package (Gu *et al*., 2014). The R package ‘xtable’ (Dahl, 2016) was used to generate LATEX tables in R.

### 4.8 Data availability

Phenotypes and genotypes for the CNLM population can be found in Santantonio *et al*., (2018b). A list of homeologous genes can be found in supplementary file ‘homeoGeneList.txt’. The supplementary file ‘HomeoMarkerSet.txt’ contains non-unique marker sets anchored to each homeologous gene set. Unique marker sets used can be found in ‘uniqueHomeoMarkerSet.txt’, ‘WithinMarkerSet.txt’, ‘AcrossMarkerSet.txt’ for the Homeo, Within and Across marker sets. Marker and marker interaction effect estimates and p-values for the Homeo set can be found in ‘twoWayInteractions.txt’ and ‘threeWayInteractions.txt’ for two- and threeway marker interactions, respectively. Phenotypes and genotypes used in the RIL population are included in the ‘NY8080Cal.txt’ file.

## 5 Results and Discussion

### 5.1 Rht-1

#### 5.1.1 RIL population

The markers linked to the *Rht-1B* and *Rht-1D* genes both had significant additive effects (p < 10^−10^) and explained 19.6% and 20.5% of the variation in the height of the RIL population (Supplemental Table S1). The test for a homeoallelic epistatic interaction between these *Rht-1* linked loci was also significant (p = 0.0025), but only explained 3.5% of the variance after accounting for the additive effects. Had we tested all pairwise marker interactions in this population, this test would not have passed a Bonferroni corrected significance threshold.

Effect estimates for the *Rht-1* markers and their epistatic interaction are shown in Table 2, for {0, 1} and {−1, 1} marker parameterizations, and for orientations where the marker main effects are both positive or both negative. The {0, 1} parameterization is arguably more intuitive, as effects correspond directly to differences in genotype values (Figure 3). They both contain the same information and are equivalent for prediction using ordinary least squares, but the interpretation of the {−1, 1} marker coding is less obvious because the slopes are deviations from the expected double heterozygote (assuming no dominance), which does not exists in an inbred population. The {0, 1} parameterization uses the double dwarf as the reference point, where the effects «β and *ac* are the two semi-dwarf genotypes. The tall genotype is the sum of the semi-dwarf allele effects plus the deviation coefficient, *δ*, which corresponds to (*αα*)*_BC_*.

**Figure 3.**
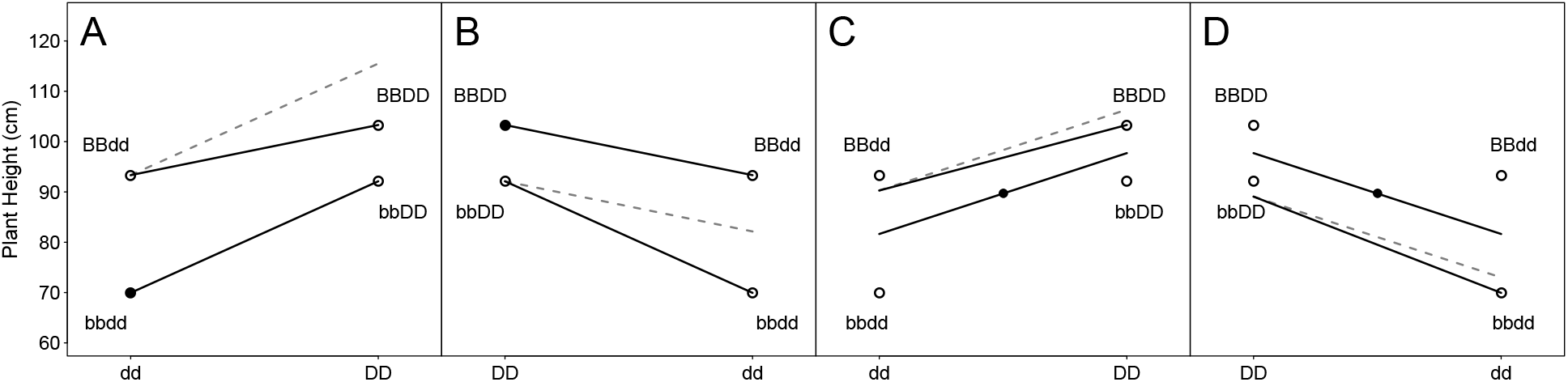
Epistasis plot of effects for *Rht-1B* and *Rht-1D* linked markers on plant height in 158 RIL lines derived from NY91017-8080 × Caledonia. The filled circles indicate the intercept (i.e. reference point) for each model parameterization while open circles indicate genotype class means. The solid lines indicate the marker effect estimates including the interaction term, while the dotted line indicates the expectation based on the additive model. **A**) {0, 1} marker coding with positive marker effect orientation. **B**) {0, 1} marker coding with negative marker effect orientation. **C**) {−1, 1} marker coding with positive marker effect orientation, **D)** {−1, 1} marker coding with negative marker effect orientation.

**Table 2:**
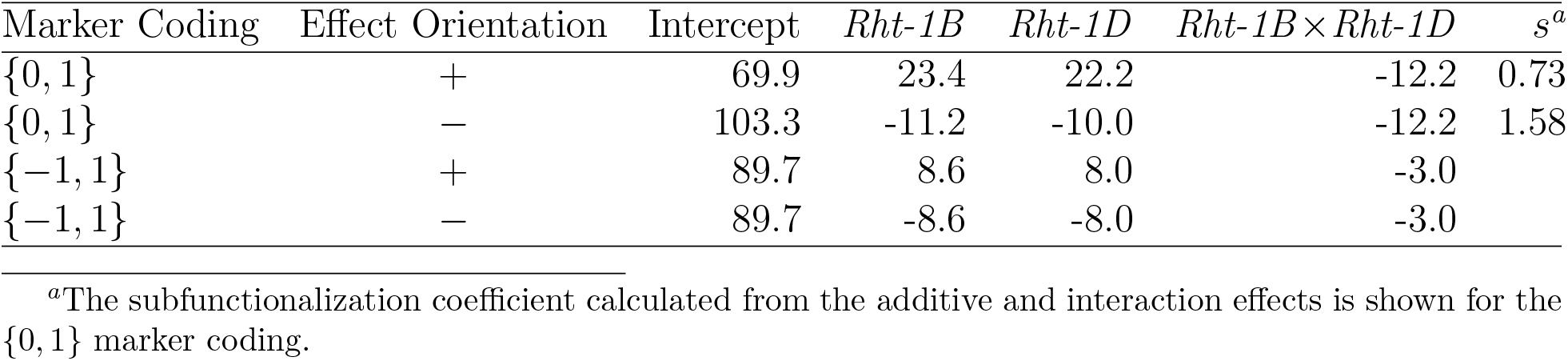
Marker and epistatic effect estimates for *Rht-1D* and *Rht-1B* linked GBS markers for plant height (cm) in 158 RIL lines derived from NY91017-8080 × Caledonia. Least squares effect estimates are for markers coded either using {0, 1} coding or {−1, 1}, and then oriented such that the two marker main effects are either both positive (+) or both negative (−).

The estimated *s* parameter of 0.73 indicates a significant degree of redundancy between the wild type *Rht-1* homeoalleles. This suggests that either the gene products maintain partial redundancy in function, or the expression of the two homeoalleles is somewhat redundant. The latter is less likely given that the two functional wild type genes have comparable additive effects relative to the double dwarf. If the two genes were expressed at different times or in different tissues based on their native subgenome, the additive effects would be likely to differ in magnitude. This demonstrates a functional change between homeoalleles that has been exploited for a specific goal, semi-dwarfism.

When the markers are oriented in the opposite direction, to indicate the GA insensitive mutant allele as opposed to the GA sensitive wildtype allele, the interpretation of the interaction effect changes. The additive effect estimates become indicators of the reduction in height by adding a GA insensitive mutant allele. The interaction effect becomes the additional height reduction from the additive expectation of having both GA insensitive mutant alleles, resulting in a *s* parameter of 1.58. The same interpretation can be made, but must be done so with care. Losing wildtype function at both alleles results in a more drastic reduction in height than expected because there is redundancy in the system. Therefore, the *s* parameter is most easily interpreted when the functional direction of the alleles is known. Simply put, when function is added on top of function, little is gained, but when all function is removed, catastrophe ensues.

#### 5.1.2 CNLM population

For the CNLM population, the markers with the lowest p-values associated with plant height on the short arms of 4B and 4D did not show a significant interaction with their respective assigned homeologous marker in homeologous sets H4.16516 and H4.23244, respectively. A new homeologous marker set, CNLM_Rhtl, was constructed with the SNPs on 4BS and 4DS with the lowest p-values mentioned above. The additive effects of markers S4B_PART1_38624956 and S4D_PART1_10982050 had p-values of 5.5 × 10^−4^ and 3.7 × 10^−8^, respectively, while the interaction had a p-value of 0.015. This set was oriented in the same direction as the RIL_Rhtl set using the LAVHAE orientation method. While the magnitude of these effects was reduced (7.13, 7.09 and −4.56 for the 4D, 4B and 4B×4D effects respectively), the CNLM_Rhtl set had a *s* parameter value of 0.68, similar to that of RIL_Rhtl. Had this set alone been tested, we would have concluded that this was a significant homeologous interaction.

To verify these results, we genotyped 1,259 individuals of the CNLM population with two ‘perfect’ markers designed to track the *Rht-1B* and *Rht-1D* alleles (Ellis *et al*., 2002). When correcting for population structure, effect estimates were 1.66 (p = 3.3 × 10^−2^), 1.93 (p < 2 × 10^−16^) and −1.02 (p = 6.4 × 10^−6^) for the *Rht-1B Rht-1B* and *Rht-1B* × *Rht-1B* terms, respectively, resulting in an *s* value of 0.71. The relatively high p-value for the *Rht-1B* is likely due to correction for population structure, where the *Rht-1Db* dwarfing allele is the predominant source of semi-dwarfism in the breeding population (Supplementary Table S3). Ignoring population structure produced p-values of p < 10”^19^ for both additive effects and p = 5.7 × 10^−5^ for the interaction.

### 5.2 Significant homeoallelic interactions

The absence of one genotype class in 7,912 interaction terms resulted in 20,641 testable interaction effects out of 28,553 total interaction terms. A trait-wise Bonferroni significance threshold of 0.05/20, 641 = 2.4 × 10^−6^ was therefore used to determine which interaction effects had a significant effect on the phenotype. Few homeoallelic interactions were significant at the trait-wise Bonferroni cutoff (Figure 4). Significant homeoallelic interactions for PH were identified between 4AL and 4DS, as well as 4BL and 4DL. Both of these locations were likely too far away from the *Rht-1* alleles to be tagging these genes directly, but they may be regulatory sites for these genes. Another set of interacting sites between the homeologous chromosome arms 3AS, 3BS and 3DS was also identified for PH, but the additive effects were not significant. Two interacting regions on homeolog 1, between IAS and IDS and between 1AL and 1DL, and three interacting regions on homeolog 5 also appeared to be influencing HD. One region on the distal end of homeolog 7 affected both HD and TW, with significant two-way and three-way interactions. Although they were tagged with different marker sets for the two traits, these epistatic regions appeared to co-localize within 2 Mbp.

**Figure 4.**
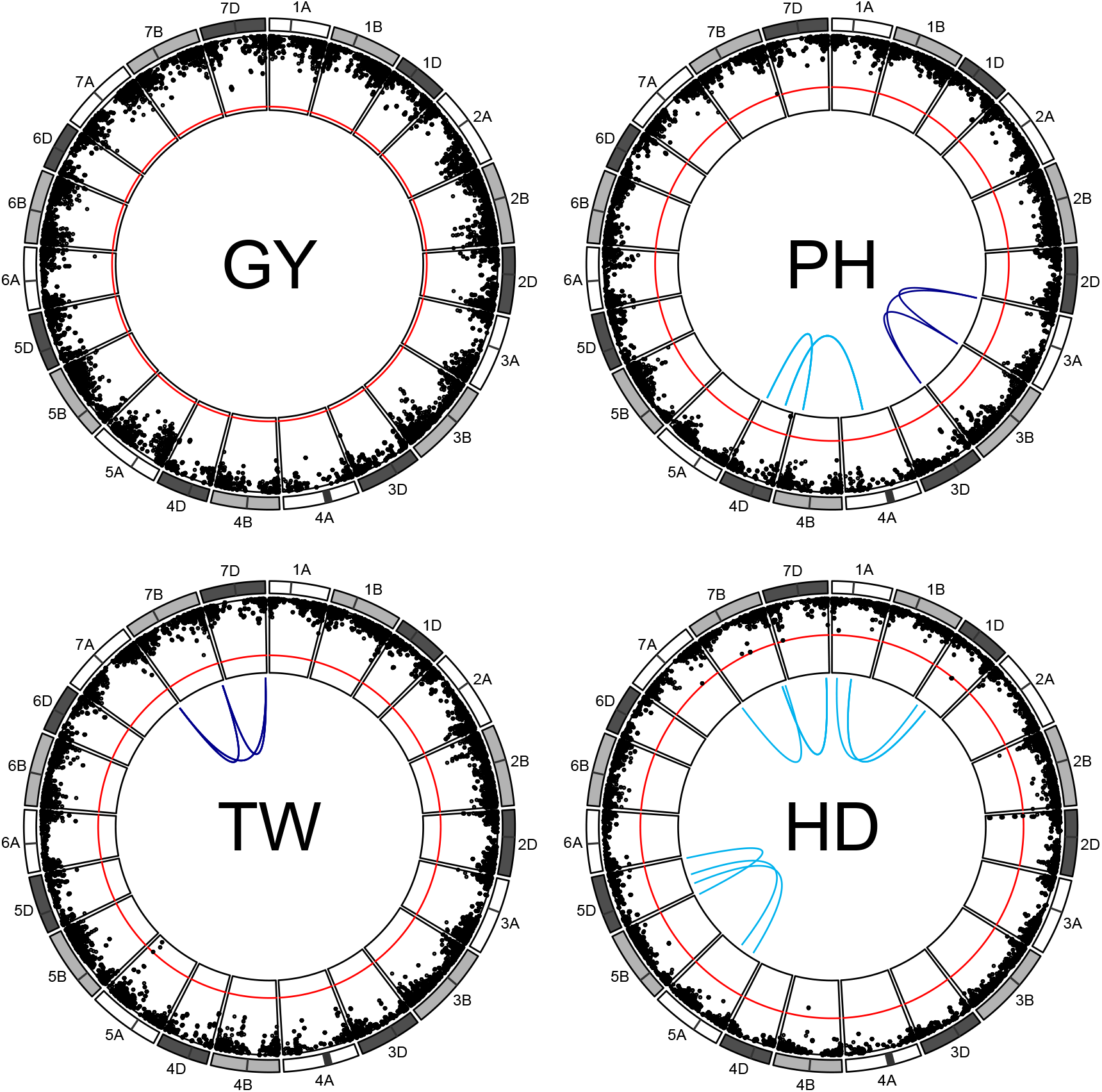
Manhattan plot of homeoallelic marker sets for each of the 21 chromosomes of wheat. The red line indicates a trait wise Bonferroni significance threshold for additive effects of –log_10_(6.0 × 10^−6^) = 5.2. Light blue lines indicate significant two-way homeoallelic marker interactions that exceeded a Bonferroni threshold for all testable interaction effects –log_10_(2.4 × 10^−6^) = 5.6. Dark blue lines indicate significant 3-way homeoallelic marker interactions that exceeded the same Bonferroni threshold.

No significant additive or interaction effects were detected for GY, highlighting the highly polygenic nature of the grain yield trait. In several cases, one of the additive effects was significant but the other was not, and it is not clear if this is influencing the detection of interactions. It may be that the significant marker is simply in higher LD with the functional mutation conditional on the presence of the other marker, allowing the interaction to pick up the additional signal from the functional mutation (Wood *et al*., 2014). However, if this were the case, the interaction would be expected to be in the same direction as the additive effect, which was not generally observed.

We did not detect an interaction between the two significant additive regions on 2B and 2D for the HD trait. While these two markers were not grouped as a homeologous set, they were tested as such based on their proximity to the well described Photoperiod-1 genes, *Ppd-Bl* and *Ppd-Dl*, on chromosomes 2B and 2D respectively. These genes are known to influence photoperiod sensitivity, and therefore transition to flowering and heading date (Welsh J.R. and R.D., 1973; Law *et al*., 1978; Scarth and Law, 1983). Certain allele pairs at these genes have been shown to exhibit a high degree of epistasis (Poland, 2018, personal communication) in a bi-parental family. It is unclear why no interaction was observed in this population.

Jiang *et al*., (2017) also investigated the presence of homeologous interactions, but found little evidence in a large population of hybrid wheat. They did not attempt to tag homeologous loci, but instead considered interactions across any markers on homeologous chromosomes to be syntenic. Interactions at homologous and non-homeologous loci may have largely outweighed interactions across homeologous loci in that population, given it was constructed from highly divergent parents and that progeny were not inbred. Additionally, they tested all pairwise marker combinations, resulting in a strict significance threshold that may have missed small effect homeologous interactions.

Homeologous interactions make up relatively few of the potential two-way interactions within an allopolyploid genome. Given a subgenome with *k* genes and alloploidy level *p* (i.e. the number of subgenomes), there are 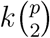 two-way homeologous interactions versus 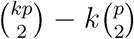 potential two-way non-homeologous gene interactions. For a subgenome size of 30,000 genes, this represents 0.02% and 0.006% of the possible two-way gene interactions for an allotetraploid and an allohexaploid, respectively. That said, homeoallelic interactions should be far more likely to have a true biological interaction than random pairs of genes because they should belong to the same or similar biochemical pathways.

### 5.3 Estimates of the subfunctionalization coefficient

There were few cases where at least two additive effects and their corresponding interaction effect were all significantly different from zero. This may be due to the difficulty of assigning functional homeologous gene sets using single SNPs, as well as a lack of statistical power owing to low minor allele frequencies (Hill *et al*., 2008). The lack of a large number of significant interactions is not surprising given that allele frequencies near 0.5 are uncommon in both natural and breeding populations.

To determine whether more homeologous marker sets were displaying a pattern indicative of subfunctionalization than would be expected by chance, marker sets where both additive and two-way interaction effects were significant at a threshold of *α* = 0.05 were examined (Table 3). The expected number of two-way marker sets with significant additive and interaction effects is about 11 (i.e. 4 traits × 22,411 two-way interactions × 0.05^3^), assuming independence of loci and true additive and interaction effects of zero. Only the Homeo and Across marker sets had significantly more than expected. When broken down by trait, these appeared to be driven by interactions for PH and TW in the Homeo set (Supplementary Table S16). The homeologous marker set had a larger proportion of *s* coefficients estimated between 0.5 and 1 relative to the strictly additive simulated phenotypes as well as the other non-homeologous marker sets, suggesting that homeologous loci exhibit a pattern indicative of subfunctionalization more so than other marker sets tested. The Across set showed the highest proportion of *s* < 0.5, suggestive of gene pathway interference. Because the power to detect significant effects diminishes as more tests are accomplished, it may be prudent to look at global trends between homeologous additive effects and their interactions, regardless of statistical significance.

**Table 3:** Estimates of *s* coefficients for marker sets where both additive and the two-way interaction effects were significant at *p* < 0.05, combined for all 4 traits. The expected number of non-zero additive and two-way interactions effects based on a 0.05 significance threshold by chance is 11 (i.e. 4 traits × 22,411 two-way interactions × 0.05^3^). Coefficients have been grouped by categories related to the potential mode of epistasis, where *s* < 0.5 indicates a highly negative interaction, 0.5 ≤ *s* < 1 a less-than-additive interaction may be indicative of subfunctionalization for homeologous genes, and *s* > 1 which indicates positive, or greater-than-additive, epistasis. Three marker sets are shown, either across all homeologous loci (Homeo), sampled sets within (Within) and across (Across) non-syntenic subgenome regions. An additional phenotype was simulated to contain additive only phenotypes to contain no epistasis, and fit with the Homeo marker set (Simulated Additive).

| Marker Set | $s < 0.5$ | $0.5 \leq s < 1$ | $s > 1$ | Total<sup>a</sup> |
| --- | --- | --- | --- | --- |
| Homeo | 8 | 14 | 8 | 30*** |
| Simulated Additive | 1 | 1 | 4 | 6 |
| Across | 9 | 7 | 1 | 17* |
| Within | 6 | 3 | 4 | 13 |

<sup>a</sup>\*, \*\*, \*\*\* indicate significantly greater than the expected number of significant sets at $p = 0.05$ , 0.01 and $10^{-6}$ based the binomial distribution with 89,644 trials and a probability of 0.05<sup>3</sup>.

### 5.4 Evidence of subfunctionalization

A strong negative relationship between additive and interaction effects was observed when using the {0, 1} marker parameterization (Figure 5A). This negative relationship was also observed in the phenotypes simulated to be strictly additive (Supplemental Figure S3). The multicollinearity of the additive and epistatic predictors at least partially drives this relationship, where positively correlated additive and epistatic predictors will tend to have effect estimates in opposing directions.

**Figure 5.**
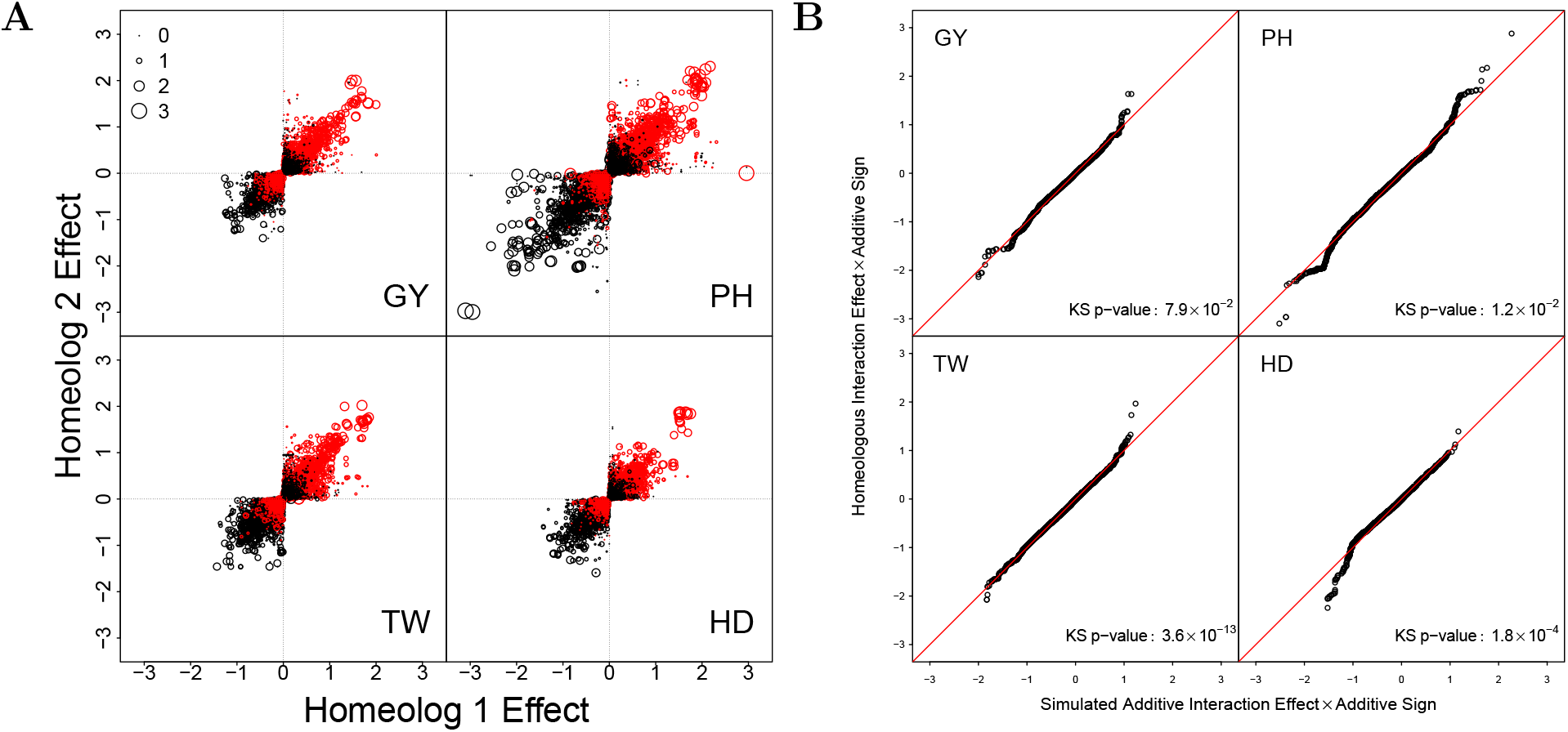
**A)** LAVHAE oriented homeologous marker pair additive effects for four traits, GY, PH, TW and HD. Point size represents the magnitude of the two-way homeologous interaction effect while color denotes the direction of the interaction effect, where black is positive and red is negative. **B)** Quantile quantile plot of the ordered estimated homeologous interaction effects plotted against those from a simulated phenotype sampled to obtain no epistatic interactions. Interaction effects have been multiplied by the effect sign of the corresponding additive effects to emphasize the relationship between the additive and interaction effects. The lower left quadrant indicates a less-than-additive interaction, whereas the upper right quadrant indicates a greater-than-additive interaction. The p-value from a Kolmorgorov-Smirnov (KS) test is reported to test if the distributions of actual and simulated interaction effect estimates are the same. A deviation below the line on the bottom left of each graph (i.e. a low dropping tail) should indicate a less-than-additive epistatic pattern of subfunctionalization, whereas a deviation above the line in the upper right (i.e. a high rising head) should indicate a greater-than-additive epistasis pattern of homeologous over dominance.

To determine if the interaction effects were greater in magnitude than expected by chance, the ordered interaction effects from the true and simulated phenotypes were plotted against one another to form a quantile-quantile plot (Figure 5B). The interaction effects were multiplied by the sign of the corresponding additive effects to highlight the direction of interaction effect relative to the additive effect. Interaction effect distributions were significantly different between the observed and strictly additive simulated data as determined by the Kolmogorov-Smirnov test (KS; p < 0.05) for all traits except GY.

HD showed a pattern consistent with a subfunctionalization model, with a low dropping tail for interaction effects in the opposite direction than that of the corresponding additive effects. This indicates that the less-than-additive effects of some estimated interactions are greater than expected by additivity alone. PH showed some evidence of this pattern, but also demonstrated a greater-than-additive effect for positively related interaction effects. The LAVHAE orientation scheme may have selected the wrong marker coding for those marker sets, resulting in a *s* parameter greater than 1, or there are true greater-than-additive interaction responses for positive effect alleles. Greater than additive responses would be indicative of overdominance across homeologous loci. GY and TW showed little evidence of the less-than-additive pattern, yet TW did show this trend when the HTEV marker orientation was used (Supplemental Figures S5 and S6). These relationships were more pronounced when the markers were permuted to remove LD before simulating the data (Supplemental Figure S4). High LD between homeologous marker sets may result in dampening of the epistatic signal due to unbalanced or missing genotype classes.

These findings are further supported by comparing the homeologous interactions to the Within and Across interaction effect estimates. The Homeo marker set showed more severe less-than-additive epistasis than both Within and Across for HD but not the other traits (Supplementary Figures S7 and S8). The Within set had more severe less-than-additive interaction effects than the Homeo set for TW (Supplemental Figure S7), and the Across had more severe less-than-additive effects for PH (Supplemental Figure S8). Large or moderate effect negative epistasis is expected across subgenomes in allopolyploids, but it is unclear why this was also observed for the Within marker set for TW.

### 5.5 Homeologous model fit

Comparing variance component estimates across different unstructured covariance matrices can be misleading as variance components can be scaled by pulling a constant out of the covariance matrix. Additionally, variance partitioning is only reliable when the covariance matrices are truly independent (Vitezica *et al*., 2017; Huang and Mackay, 2016; Jiang *et al*., 2017). Therefore, we do not make an attempt to discern meaning from the variance components *per se*, and instead focus the discussion on model fit diagnostics, as well as prediction accuracy from cross validation to determine the value of the predictive information included in the model.

All epistatic models using the {−1, 1} marker parameterization provided a superior fit to the additive only model based on Akaike’s Information Criterion (AIC) for all traits (Supplementary Table S4). These results were confirmed by a likelihood ratio test to determine if the epistatic variance component was zero for all traits. With the exception of the GY trait, all of the epistatic models using the {0, 1} marker parameterization also had non-zero variance components (Supplementary Table S5), but did not result in a better fit for any models or traits. The LAVHAE method outperformed all other marker orientation schemes (Supplementary Tables S6, S7, S8 and S9). The Pairwise, Within and Across epistatic models outperformed the Homeo marker interaction set for all traits. This may be due to poor assignment of homeologous sets, or relatively fewer identifiable interactions and is discussed later.

### 5.6 Genomic prediction

All epistatic models resulted in higher prediction accuracies for all traits other than GY, where only marginal increases were seen for certain marker interaction sets and parameterizations (Table 4). The {−1, 1} marker coding resulted in higher prediction accuracies with a mean increase of 0.045 over the {0, 1} coding, and ranged from 0.007 to 0.084 higher accuracy. This increase may be due to choosing the wrong orientation using the {0, 1} marker coding effects. While these two codings are equivalent for prediction when marker effects are fixed, this is not the case for the mixed model genomic prediction environment (Martini *et al*., 2017, 2018). The discrepancy lies in shrinkage of interaction effects, where the {0, 1} marker coding should result in greater shrinkage than the {−1, 1} marker coding. This can be seen from a simple example with one observation of each genotypic class in {*bbcc, bbCC, BBcc, BBCC*}. The {−1, 1} coding would have an interaction predictor of {1, −1, −1, 1}, whereas the {0, 1} coding would have an interaction predictor of {0, 0, 0, 1}. This results in different numbers of observations per interaction class, with the {0, 1} coding contrasting 3 and 1, verses 2 and 2 for the {−1, 1} coding. Therefore the shrinkage of the {0, 1} coding should be greater than for the {−1, 1} coding. Martini *et al*., (2017), also noted that the {−1, 1} marker coding has a 50% chance of choosing the wrong marker orientation if chosen at random, whereas the {0, 1} marker coding has a 75% chance of being the wrong marker orientation.

**Table 4:** Prediction accuracies of whole genome Additive and Pairwise epistasis, along with the Homeo, Within and Across genome marker sets for both {−1, 1} and {0, 1} marker coding using the LAVHAE marker orientation.

| LAVHAE | Additive | Pairwise | Homeo<sub>-11</sub> | Homeo<sub>01</sub> | Within<sub>-11</sub> | Within<sub>01</sub> | Across<sub>-11</sub> | Across<sub>01</sub> |
| --- | --- | --- | --- | --- | --- | --- | --- | --- |
| GY | 0.601<sup>a</sup> | 0.604 | 0.606 (167%)<sup>b</sup> | 0.599 (-67%) | 0.627 (867%) | 0.600 (-33%) | 0.630 (967%) | 0.604 (100%) |
| PH | 0.559 | 0.637 | 0.606 (60%) | 0.580 (27%) | 0.652 (119%) | 0.570 (14%) | 0.650 (117%) | 0.584 (32%) |
| TW | 0.515 | 0.576 | 0.560 (74%) | 0.516 (2%) | 0.596 (133%) | 0.514 (-2%) | 0.581 (108%) | 0.525 (16%) |
| HD | 0.664 | 0.712 | 0.692 (58%) | 0.682 (38%) | 0.710 (96%) | 0.674 (21%) | 0.722 (121%) | 0.682 (38%) |

<sup>a</sup>Mean Pearson correlation between predicted and observed genetic values across 10 random 5-fold cross-validation replications.
<sup>b</sup>The percentage of the non additive genetic predictability as relative to the the Pairwise model is shown in parentheses (equation 4).

The LAVHAE marker orientation scheme was superior for prediction of all traits and marker sets for the {−1, 1} coding, but had little effect on the {0, 1} marker coding (Supplemental Tables S12, S13 and S14). This suggests that information can be gained from orienting markers relative to one another, however, it is still unlcear what strategy should be used to orient pairs of markers. In this report, marker additive effects were forced to be either all positive or all negative to model the homeologous subfunctionalization hypothesis, but there may be more biologically relevant orientations not explored here. Martini *et al*., (2017) used a categorical interaction that included a predictor for each pairwise genotype. That model was shown to be less predictive than the {−1, 1} multiplicative model, perhaps due to more linearly dependent predictors assumed to have non-zero effects. Feature selection may be useful for selecting the most informative interactions from this population of linearly dependent predictors. How an optimal set of orientations might be obtained without losing biological meaning of the orientation warrants further investigation.

The proportion of non-additive genetic signal attributable to homeologous gene interaction was determined by taking the ratio of the percent increase in prediction accuracy of the Homeo, Within or Across prediction models from the additive model to the increase in prediction accuracy due to all pairwise interactions (equation 4). All three marker sets resulted in higher genomic prediction accuracy than the additive only GBLUP model (*G*) when the {−1, 1} marker coding was used. The homeologous marker interaction set explained between 58% and 167% of the additional genetic signal from the additive model. This result supports the idea that homeologous interactions are an important feature in the wheat genome. Conversely, Within and Across epistatic marker sets always resulted in a higher increase in genomic prediction accuracy relative to the Homeo marker set for all traits. This may suggest that the homeologous marker interactions are the least important relative to other epistatic interactions within and across the subgenomes, but could also be due to the paucity of these interactions relative to all possible two-way interactions, as previously discussed.

Another explanation might be provided by the relatively higher degree of LD across Homeo marker sets than found for the Within or Across marker sets. Homeologous marker sets were selected next to one another along syntenic regions of homeologous chromosome, and more often shared two of the three homeoallelic markers (Supplemental Figures S13 and S14). The Within and Across sets appear to have sampled the entire genome better than selecting only homeologous loci, as they track more unique pairs of genomic regions. Two additional samples of each Within and Across sets were showed very similar outcomes to the samples shown here (see Supplementary Tables S10, S11 and S15).

### 5.7 Homeologous LD

The superiority of the Within and Across genomic prediction models to the Homeo genomic prediction model may indicate that homeologous interactions are relatively less important than other sets of interacting loci. However, homeologous marker sets had a much higher tendency to be co-inherited together, as seen by relatively higher standardized LD values, D’ (Lewontin, 1964), than observed for either Within (KS test p-value = 1.1 × 10^−6^) or Across (KS test p-value = 2.3 × 10^−13^) marker sets (Figure 6). The greater fixation of allele pairs at homeologous regions may explain the lack of increased prediction accuracy of the Homeo marker set, but this may not diminish the importance of homeologous interactions. As sets of interactions are fixed within the population, the epistatic variance becomes additive (Hill *et al*., 2008). The higher degree of LD, *per* se, may indicate the importance of homeologous interactions.

**Figure 6.**
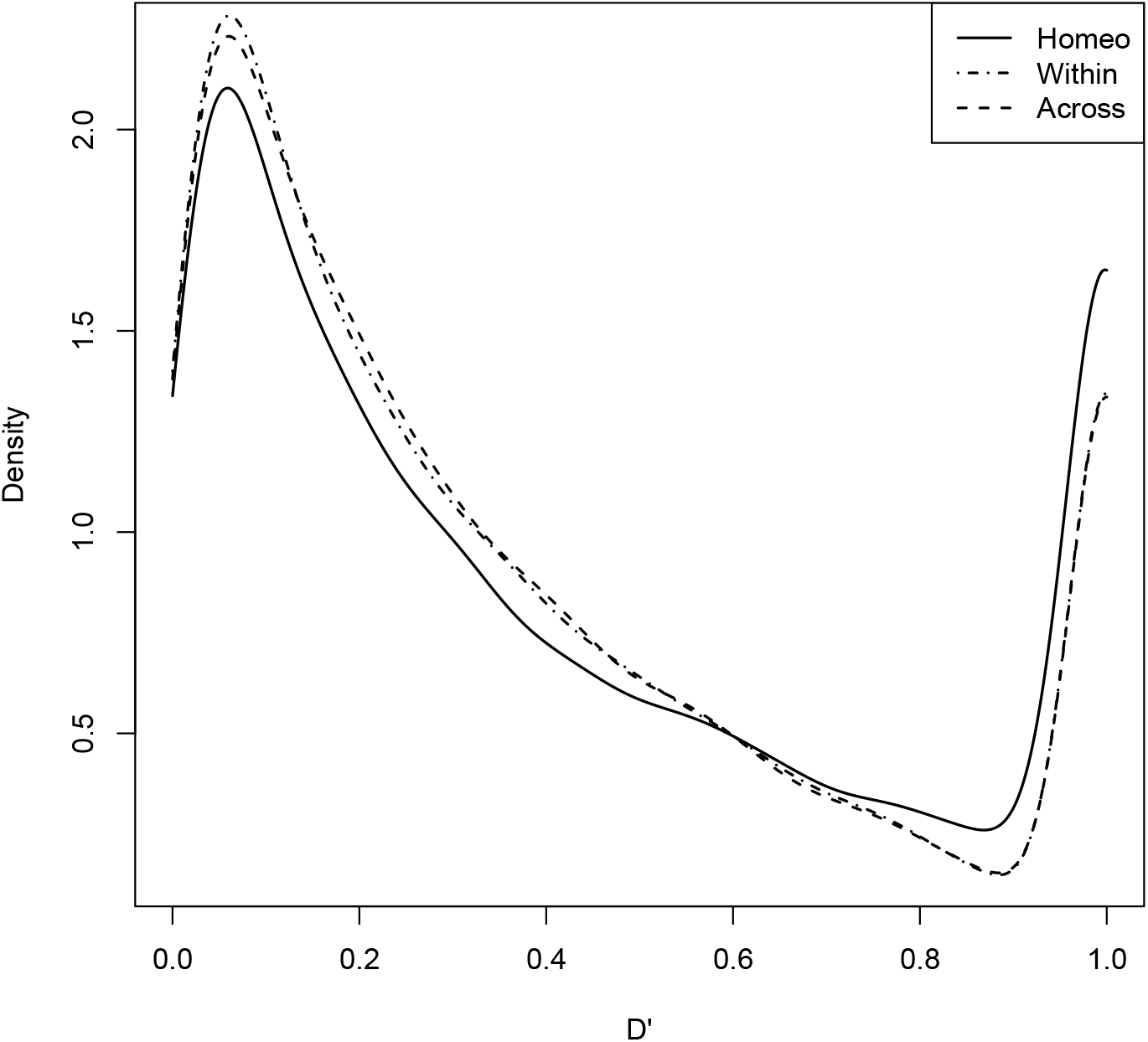
Smoothed densities of standardized D’ statistics of linkage disequilibrium for expected and observed joint allele frequencies for Homeo, Within and Across marker sets. Kolmogorov-Smirnov (KS) tests were used to determine if the distribution of LD differed between Homeo and Within (KS p-value = 1.1 × 10^−6^) or Across (KS p-value = 2.3 × 10^−13^) marker sets.

The Green Revolution dwarfing genes are an excellent example of how pairs of homeoalleles may become fixed, or develop a tendency for co-inheritance under selection. In this example, the desirable phenotype is a semi-dwarf, due to its resistance to lodging. Therefore, wildtype *Rht-1B* alleles will usually be paired with a GA-insensitive *Rht-1D* dwarfing allele, while wildtype *Rht-1D* alleles will usually be found with a GA-insensitive *Rht-1B* dwarfing allele to confer the desirable semi-dwarf phenotype. The ‘perfect’ *Rht-1* markers had a large standardized *D*′ value of 0.89, indicating that pairs of alleles were being fixed in the population.

We recognize that it is also possible that the higher degree of LD observed between homeologous marker pairs could be due to misalignment of markers to the wrong subgenome. Markers assigned to the wrong homeolog would appear in high LD simply because they are physically located near their assigned homeologous partner on the same chromosome. We used strict filtering parameters to reduce the likelihood of misalignment. This included a threshold on observed heterozygosity in the population, which could indicate alignment to more than one subgenome.

## 6 Further considerations

Wagner (2005) suggested that there are two potential drivers of less-than-additive (Eshed and Zamir, 1996) or synergistic (Segre *et al*., 2005) epistasis. These drivers are i) functional redundancy, as might be expected across homeologous loci, and ii) distributed robustness of function, in which there can be are many pathways that can acheive the same outcome. Our observation that most epistasis is not due to homeologous interactions is supported by the findings of Jannink *et al*., (2009), who found the synergistic epistasis signal in a wheat dataset to be indicative of Wagner’s distributed hypothesis, and not of the redundancy hypothesis.

It may be that there are few differences in protein function or expression across the three subgenomes, although this seems unlikely given mounting evidence that homeologous copies are differentially expressed in time, tissue and environment (Adams *et al*., 2003; Liu and Adams, 2007; Liu *et al*., 2011; Chaudhary *et al*., 2009; Pfeifer *et al*., 2014; Liu *et al*., 2015; Zhang *et al*., 2016; Mutti *et al*., 2017). We were unable to assign homeologous pairs to all genes within the genome, suggesting that many of these potential sites for interacting loci were lost during polyploidization. Rapid loss of genetic material due to genome shock (McClintock, 1984) is common in newly synthesized allopolyploids (Comai *et al*., 2003; Chen and Ni, 2006), as has been shown in synthetic allopolyploid wheat (Ozkan *et al*., 2001; Kashkush *et al*., 2002). Other interacting loci may have undergone epigenetic (Comai, 2000; Lee and Chen, 2001; Comai *et al*., 2003) or transposon induced silencing of one or more homeoalleles (Kashkush *et al*., 2003; Wang *et al*., 2004).

The large portions of duplicated genes retainment across subgenomes suggests there is a benefit to their maintenance. Duplicate copies may be important contributors to differential genotype performance in contrasting environments. Unfortunately, the CNLM dataset lacks sufficient genotype by environment variation to properly ask this question (data not shown). Experiments designed to explicitly model the phenotypic effect of differential homeologous gene expression across contrasting environments will be necessary to provide a satisfactory answer.

One of the challenges of using diverse panels of individuals is that marker proximity to a functional mutation is not necessarily indicative of high LD between the two sites. Significantly older or newer marker mutations may be in weak LD with a functional mutation despite close physical proximity, at least until a genetic bottleneck brings them back into high LD, such as in a bi-parental population (Flint-Garcia *et al*., 2003; Weir, 2008). Other strategies to determine functional homeologous regions relax which sets of markers are considered homeologous. This has been accomplished by allowing pairwise relationships with all markers across entire subgenomes (Santantonio *et al*., 2018b) or on syntenic chromosome arms (Santantonio *et al*., 2018a), with mixed success. The construction of smaller haplotypes in a manner similar to Gao *et al*., (2017) may also improve functional pairing of homeologous alleles. Higher depth sequencing and advances in marker imputation may also aid in detection of homeologous epistasis.

The TILLING population developed by Krasileva *et al*., (2017) could be a useful resource for future investigation into homeoallelic gene interactions. Lines with complementary loss of function homeologous genes could be used to develop bi-parental mapping populations to test the degree of subfunctionalization with the high statistical power afforded by allele frequencies of 0.5. So called ‘synthetic’ wheat populations formed by crossing common wheat with newly synthesized allohexaploids containing durum A and B genomes coupled to an *Ae. taushii* D genome (Sorrells *et al*., 2011, e.g.), and may prove powerful for detection of interactions between the common wheat homeologs and their durum and *Ae. taushii* ancestors.

## 7 Conclusion

While much epistasis is partitioned to additive variance, it has been shown to be prevalent (Forsberg *et al*., 2017), and is important for maintaining long term selection (Carlborg *et al*., 2006; Paixão and Barton, 2016). Our results indicate that homeologous interactions contribute to the total genetic variance of the CNLM population. However, sampling interactions across non-syntenic regions was superior for all traits examined, suggesting that homeologous epistasis make up a minority of the non-additive genetic variance. The biological state of allopolyploids, along with the suggestive evidence presented here, demonstrate that there is value in further investigation of homeologous interactions.

The most important trait, GY, showed little to no evidence of homeologous subfunctionalization. This may be due to the highly polygenic nature of the trait, where essentially all functional genetic differences in the population should contribute to GY. Modern plant breeding has likely driven large effect homeologous allele pair interactions to fixation in elite wheat genotypes. The implementation of the semi-dwarf phenotype provides perhaps the most important example where fixation of specific pairs of homeoalleles resulted in the single largest increase in wheat grain production in modern agriculture.

Prediction of unobserved homeologous allele pairs may prove difficult, as it currently is in diploid hybrids. However, large populations may be use to identify beneficial homeologous combinations that may subsequently be used for selection of unobserved lines before intensive field trials are conducted.

Treating the genome as consisting of purely additive gene action assumes that genes are independent machines, whose products sum to the final value of an individual. While convenient for selection, this is almost certainly not true when we consider the molecular mechanisms of biological organisms. Instead, genes work in concert to produce an observable phenotype. To this day, breeders of allopolyploid crops have treated allopolyploids as diploids for simplicity, but we now have the technical ability to view and start to breed these organisms as the ancient immortal hybrids that they are.

## 8 Acknowledgments

Funding of this research was provided by the USDA National Needs Fellowship for N. Santantonio, in partial fulfillment of the requirements for a Ph.D in Plant Breeding and Genetics at Cornell University. The field trials comprising the phenotypic data for the CNLM population were funded in part by the Hatch Project # 149-447. Genotyping was funded by the Wheat Coordinated Agricultural Project (WheatCAP). We are grateful to Jesse Poland’s research group at Kansas State University for their contribution to genotyping of CNLM materials. We thank Gina Brown-Guedira at the USDA-ARS Plant Science Research in the Department of Crop Science at North Carolina State University for genotyping the CNLM population for the *Rht-1B* and *Rht-1D* loci. The author give special thanks the International Wheat Genome Sequencing Consortium for pre-publication access to IWGSC RefSeq vl.0. The authors would also like to acknowledge Roberto Lonzano Gonzalez for the suggestion of using the coding sequences to identify homeologous genes. Finally, we would like to acknowledge the Cornell small grains staff, particularly David Benscher and James Tanaka, who were vital in implementing, collecting and processing the materials used to build the CNLM dataset.

## A1 Appendix 1 RIL population

The population was formed from a cross between two Cornell soft winter wheat lines, NY91017-8080 and Caledonia. Caledonia contains a GA-insensitive 4D allele, *d*, and a wildtype 4B allele, *B*, while NY91017-8080 has a GA-insensitive 4B allele, *b*, and the wild type 4D allele, *D.* The population consisting of 192 individuals was planted in single row plots in Ithaca NY and measured for plant height in 2008. The population was screened for loci influencing plant height on chromosomes 4B and 4D using genotyping by sequencing (GBS) markers. The markers with the lowest p-value on the short arms of 4B and 4D were used to indicate the *Rht-1* gene in this study. Only individuals with homozygous genotype calls for both loci were included to test for epistasis. This resulted in 19 double dwarfs (*bbdd*), 51 D genome semi-dwarfs (*BBdd*), 35 B genome semi-dwarfs(*bbDD*), and 53 tall (*BBDD*), for a total of 158 individuals. It appears that the Caledonia parent plant used in the cross was heterozygous for the D genome dwarfing allele, resulting in the 1:2 segregation ratio for the *d: D* alleles, and was confirmed by the genotype call for that plant.

## A2 Appendix 2 Coding Sequence Alignment

Alignments of coding sequences was accomplished with BLAST+, allowing up to 10 alignments with an e-value cutoff of le-5. Alignments were only considered if they aligned to 80% or more of the query gene. Of the 110,790 coding sequences, 13,111 triplicate sets with one gene on each homeologous chromosome (representing 39,333 genes) were identified with no other alignments meeting the criterion. An additional 5,073 triplicates (representing 15,219 genes) were added by selecting the top 2 alignments if they were on the corresponding homeologous chromosomes. Duplicate sets were also included if there was not a third alignment to one of the three sub-genomes, adding an additional 5,612 duplicates. The coding sequences for which we did not identify homeologous genes either appeared to be singletons (24,695 coding sequences) that did not have a good alignment to a gene on a homeologous chromosome, or had many alignments across the genome making it impossible to determine with certainty which alignments were truly homeologous (20,319 coding sequences).

## A3 Appendix 3 Change of reference

Following Álvarez-Castro and Carlborg (2007), we demonstrate the change-of-reference operation simplified for inbred populations. For {0, 1} marker coding and allowing *G*_1_ to be the reference genotype, the genotypic values at a single locus can be represented as

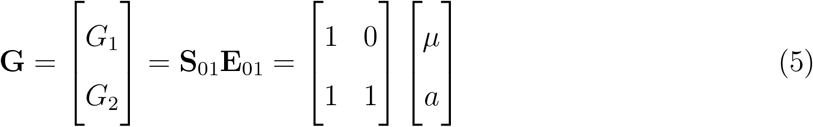

where **S**_01_ is the marker score matrix using the {0, 1} marker parameterization and **E**_01_ is the vector of expected values. For the two locus epistasis model, the four genotypic values are then

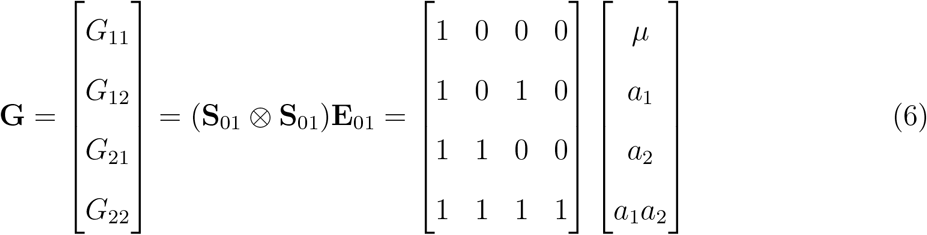

The three locus interaction is extended by

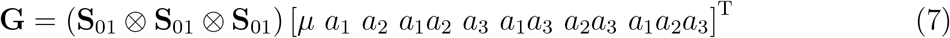

To shift from {−1, 1} coding estimates, *β*_−11_·, to {0, 1} coding estimates, *β*_01_ the following transformation exists (Álvarez-Castro and Carlborg, 2007). Let **S**_-11_ indicate the {−1, 1} marker parameterization

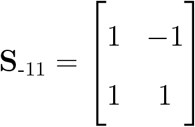

then 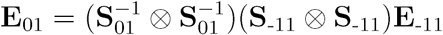.

## SI Supplementary Materials

**Table S1:** ANOVA table for *Rht-1B* and *Rht-1D* linked GBS markers and their epistatic interaction for plant height (cm) in 158 RIL lines derived from NY91017-8080 × Caledonia.

| Source | df | SS | MS | F value | $-\log_{10}(\text{p-value})$ |
| --- | --- | --- | --- | --- | --- |
| SNP36427 | 1 | 7065 | 7065 | 53.5 | 10.9 |
| SNP11172 | 1 | 7391 | 7391 | 56.0 | 11.3 |
| SNP36427:SNP11172 | 1 | 1243 | 1243 | 9.4 | 2.6 |
| Residuals | 154 | 20323 | 132 |  |  |

**Table S2:**
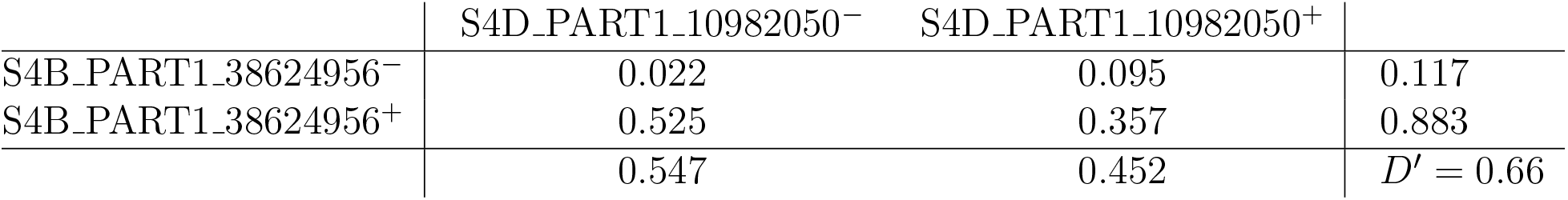
Table of genotype frequencies for the *Rht-1* linked homeologous GBS markers in the CNLM population. The + and − signs indicate the wildtype and mutant alleles, respectively. The margins indicate the marker allele frequencies.

|  | S4D_PART1_10982050<sup>-</sup> | S4D_PART1_10982050<sup>+</sup> |  |
| --- | --- | --- | --- |
| S4B_PART1_38624956<sup>-</sup> | 0.022 | 0.095 | 0.117 |
| S4B_PART1_38624956<sup>+</sup> | 0.525 | 0.357 | 0.883 |
| | 0.547 | 0.452 | $D' = 0.66$ |

**Table S3:**
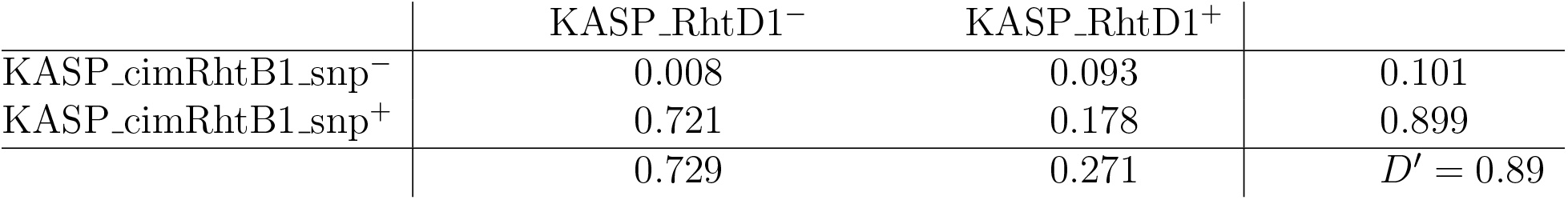
Table of genotype frequencies for the ‘perfect’ *Rht-1* markers in the CNLM population. The + and − signs indicate the wildtype and mutant alleles, respectively. The margins indicate the marker allele frequencies.

**Figure S1:**
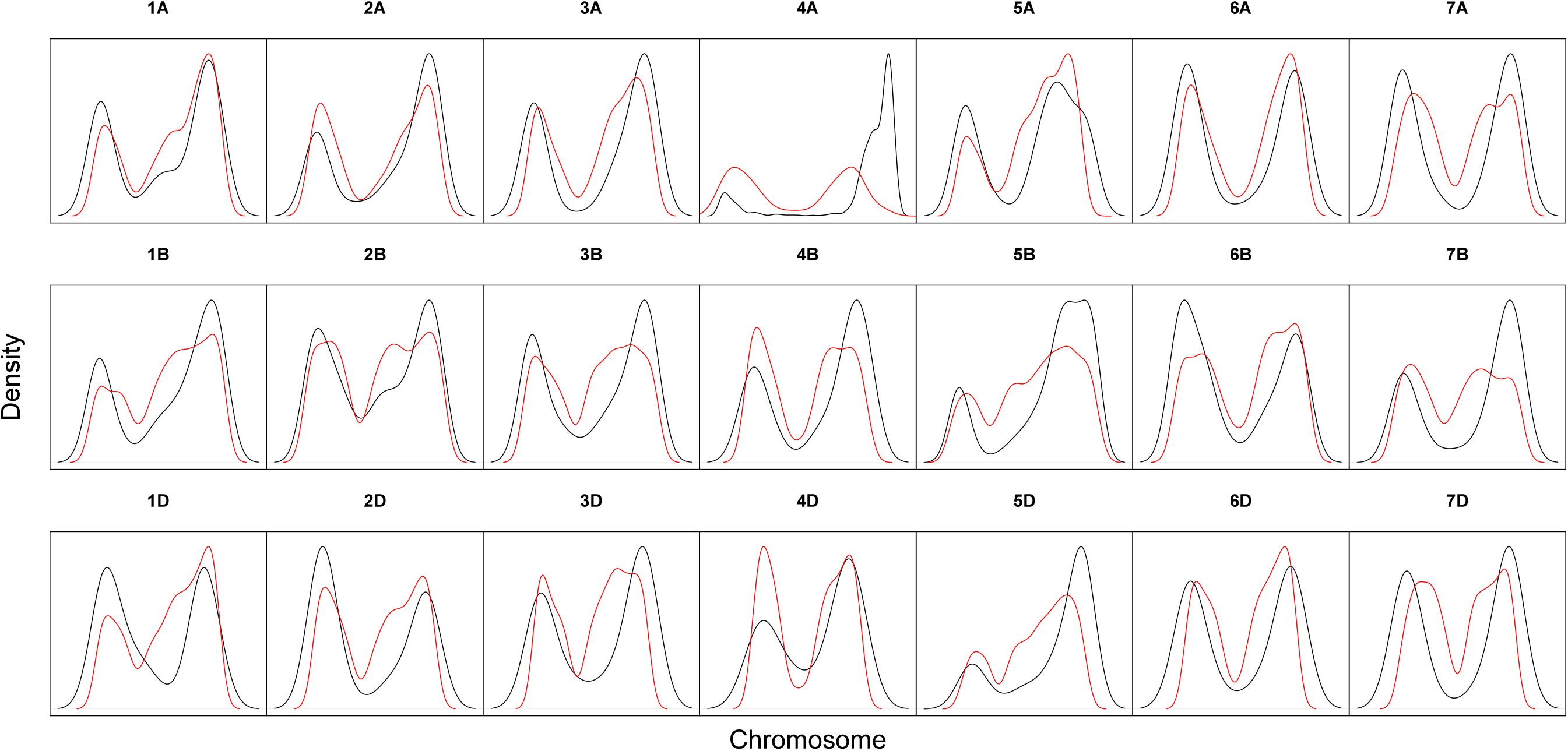
Smoothed densities of GBS markers (black) and genes (red) along the 21 wheat chromosomes in the CNLM population.

**Figure S2:**
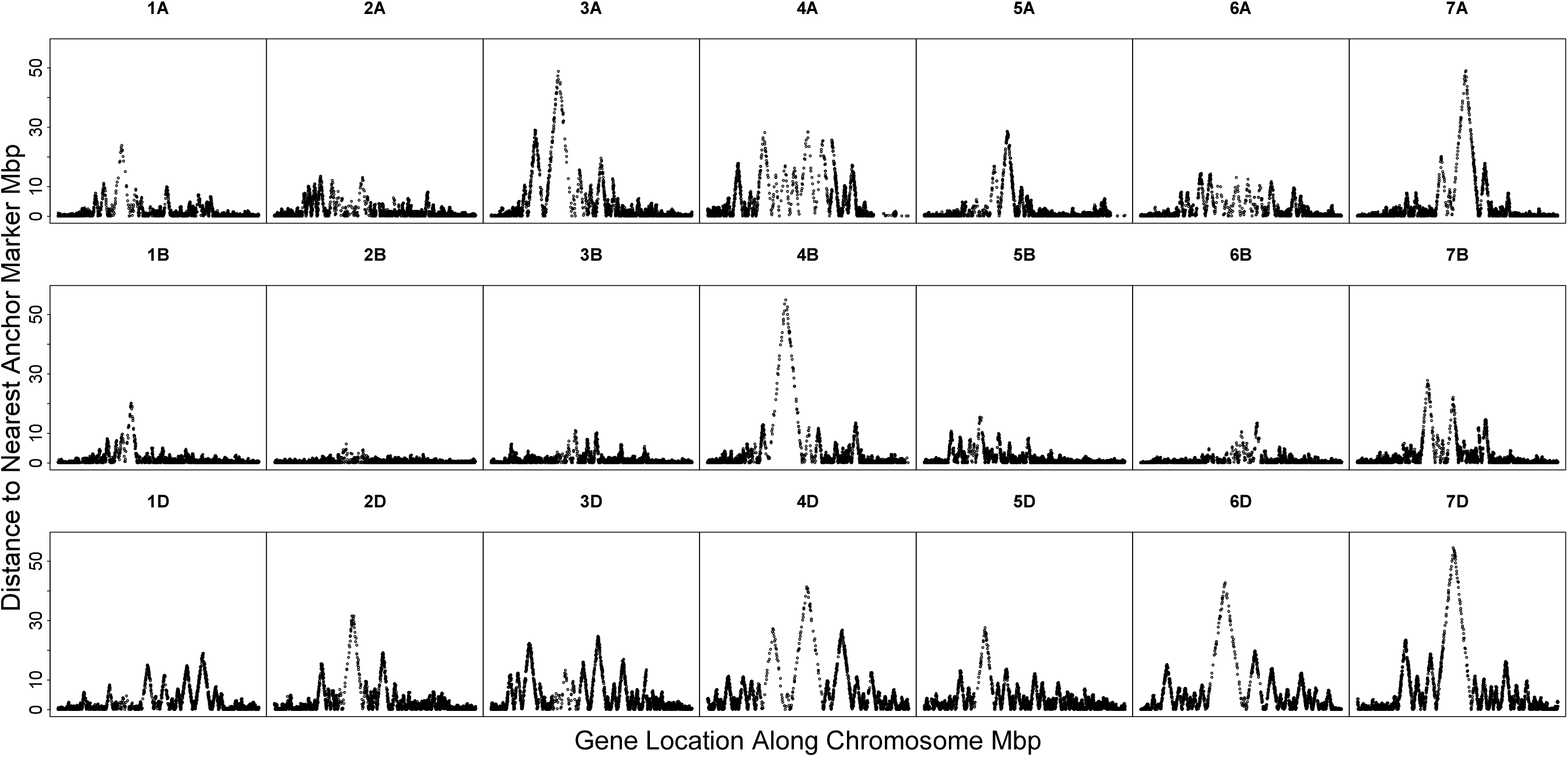
Distance of genes from their nearest GBS anchor marker along the 21 wheat chromosomes in the CNLM population.

**Figure S3:**
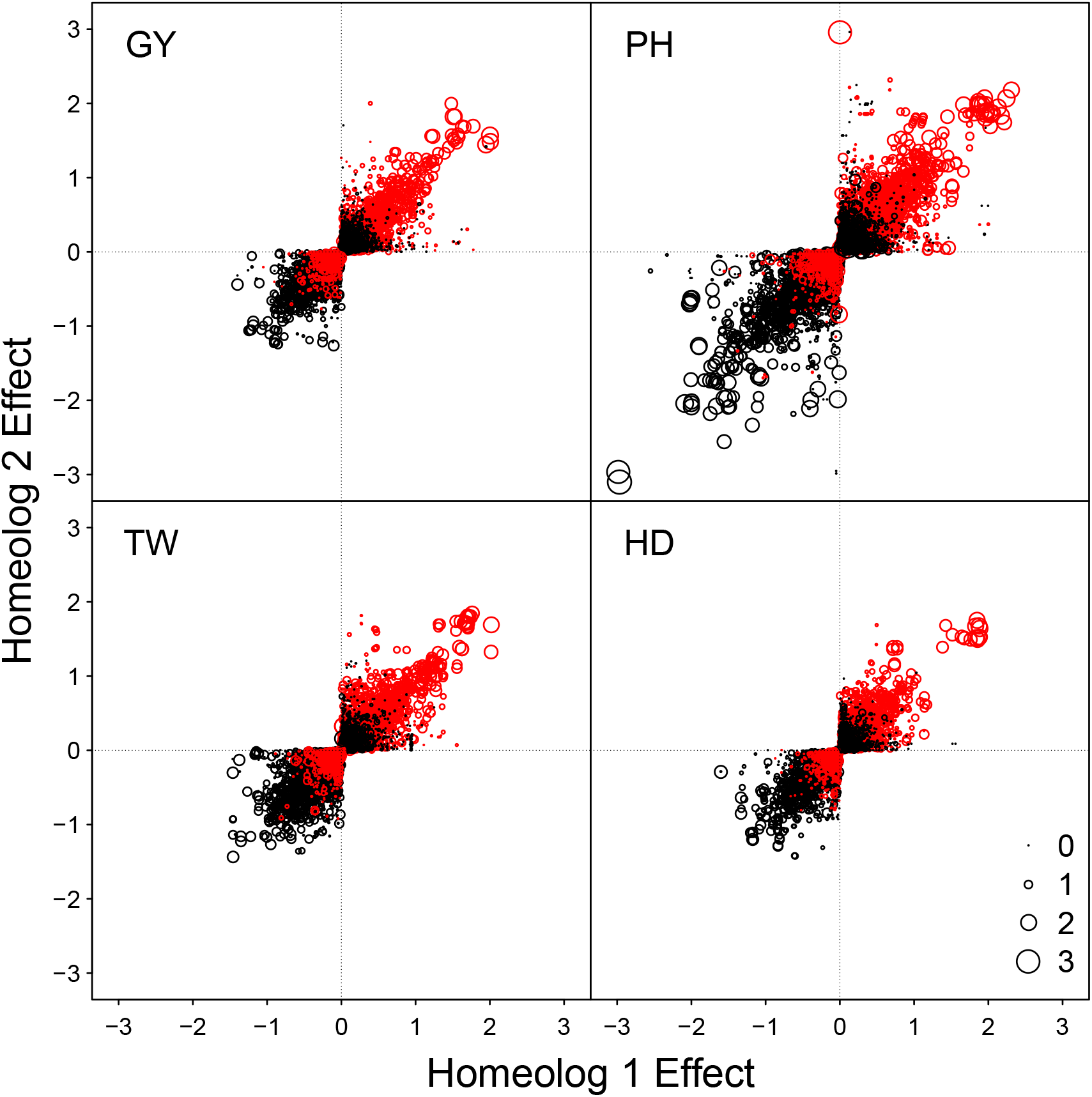
LAVHAE oriented homeologous marker pair additive effects with point size representing the magnitude of the two-way homeologous interaction effect, and the color denoting the direction of that effect where black is positive and red is negative. Four simulated phenotypes sampled to obtain no epistatic interactions, GY, PH, TW and HD, are shown.

**Figure S4:**
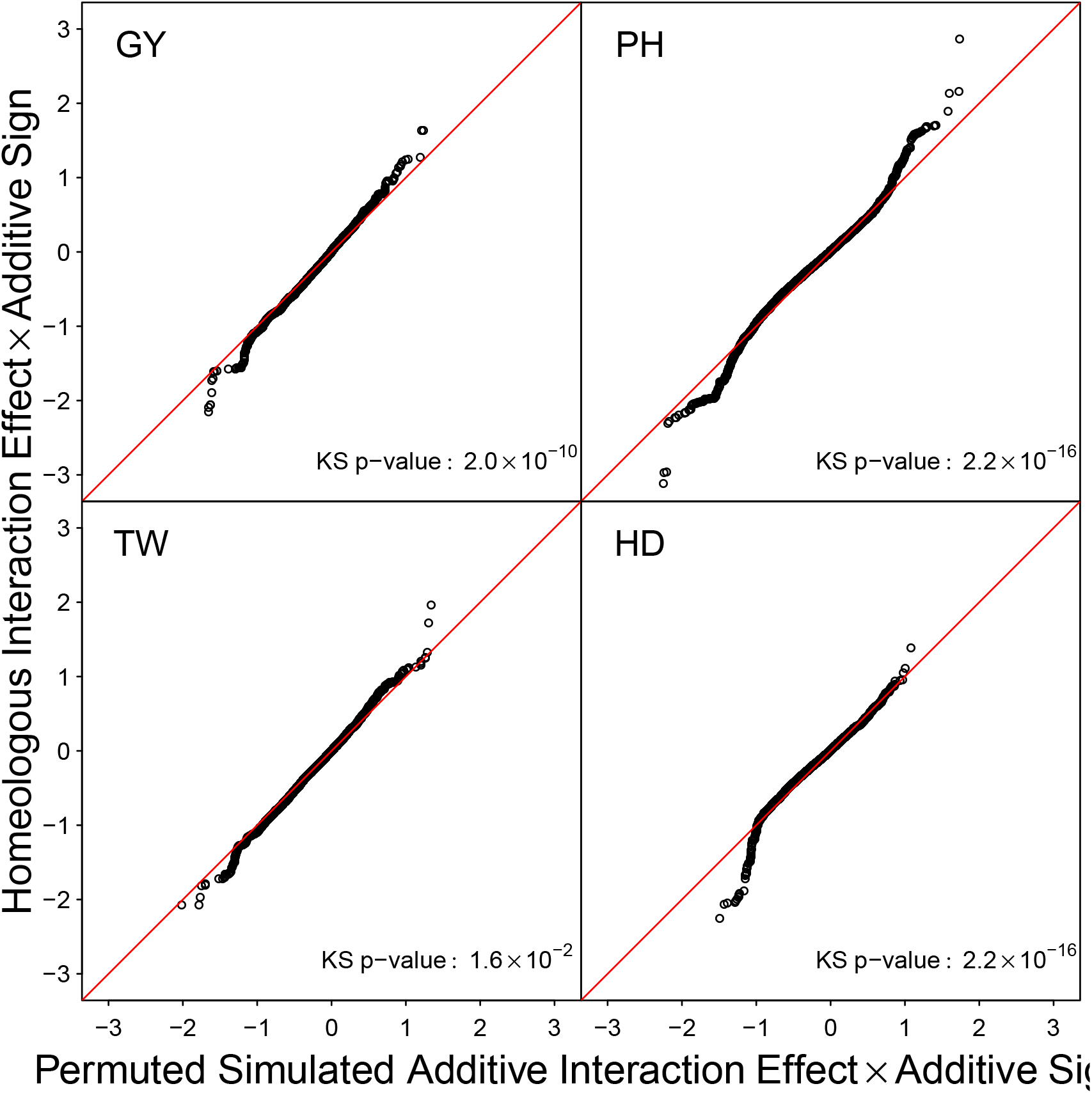
Quantile quantile plot of the ordered estimated homeologous interaction effects plotted against those from a simulated phenotype sampled to obtain no epistatic interactions using the LAVHAE marker orientation. Markers scores were permuted before simulation of the phenotype to remove LD between markers. Interaction effects have been multiplied by the effect sign of the corresponding additive effects to emphasize the relationship between the additive and interaction effects.

**Figure S5:**
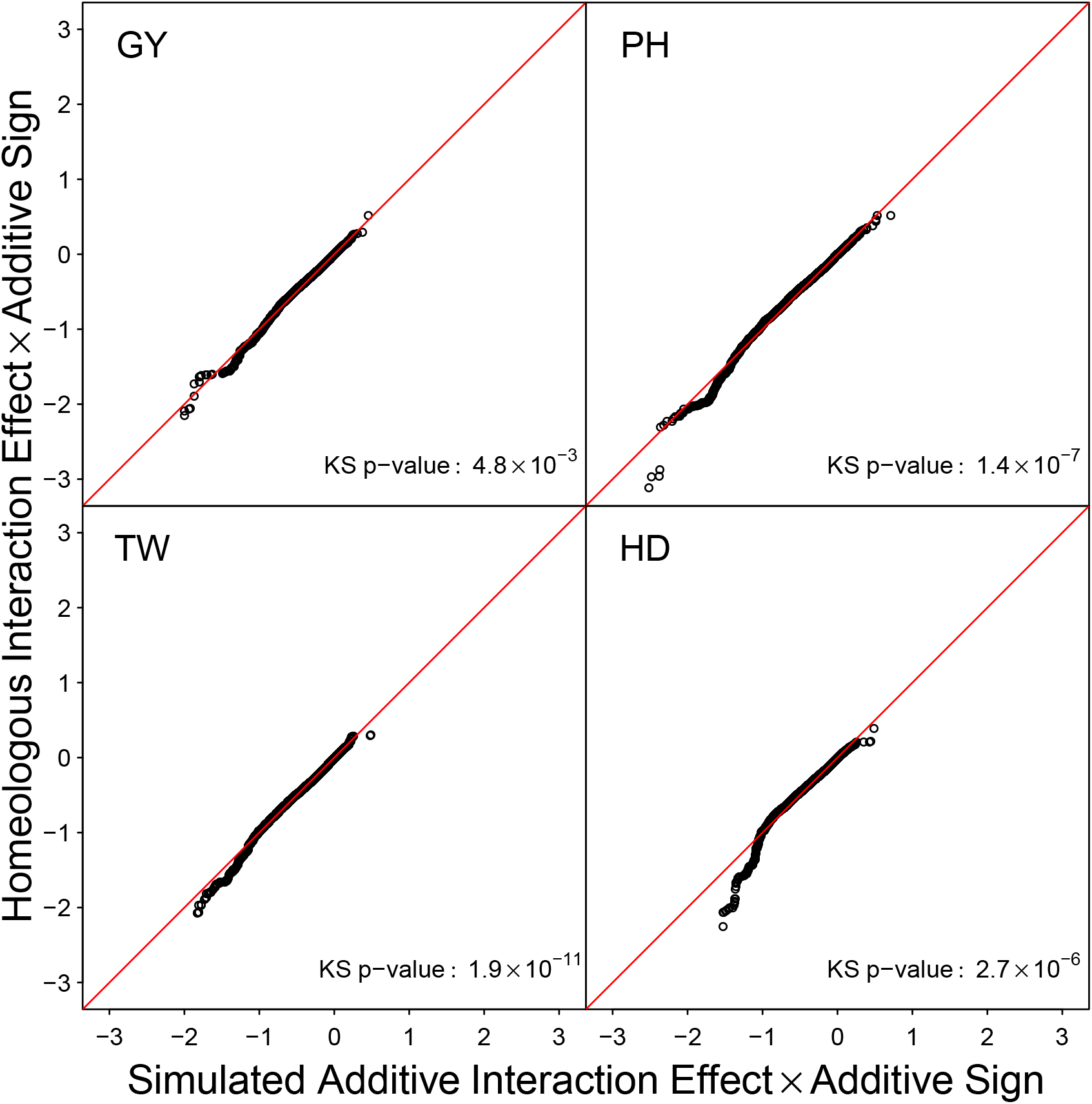
Quantile quantile plot of the ordered estimated homeologous interaction effects plotted against those from a simulated phenotype sampled to obtain no epistatic interactions using the HTEV marker orientation. Interaction effects have been multiplied by the effect sign of the corresponding additive effects to emphasize the relationship between the additive and interaction effects.

**Figure S6:**
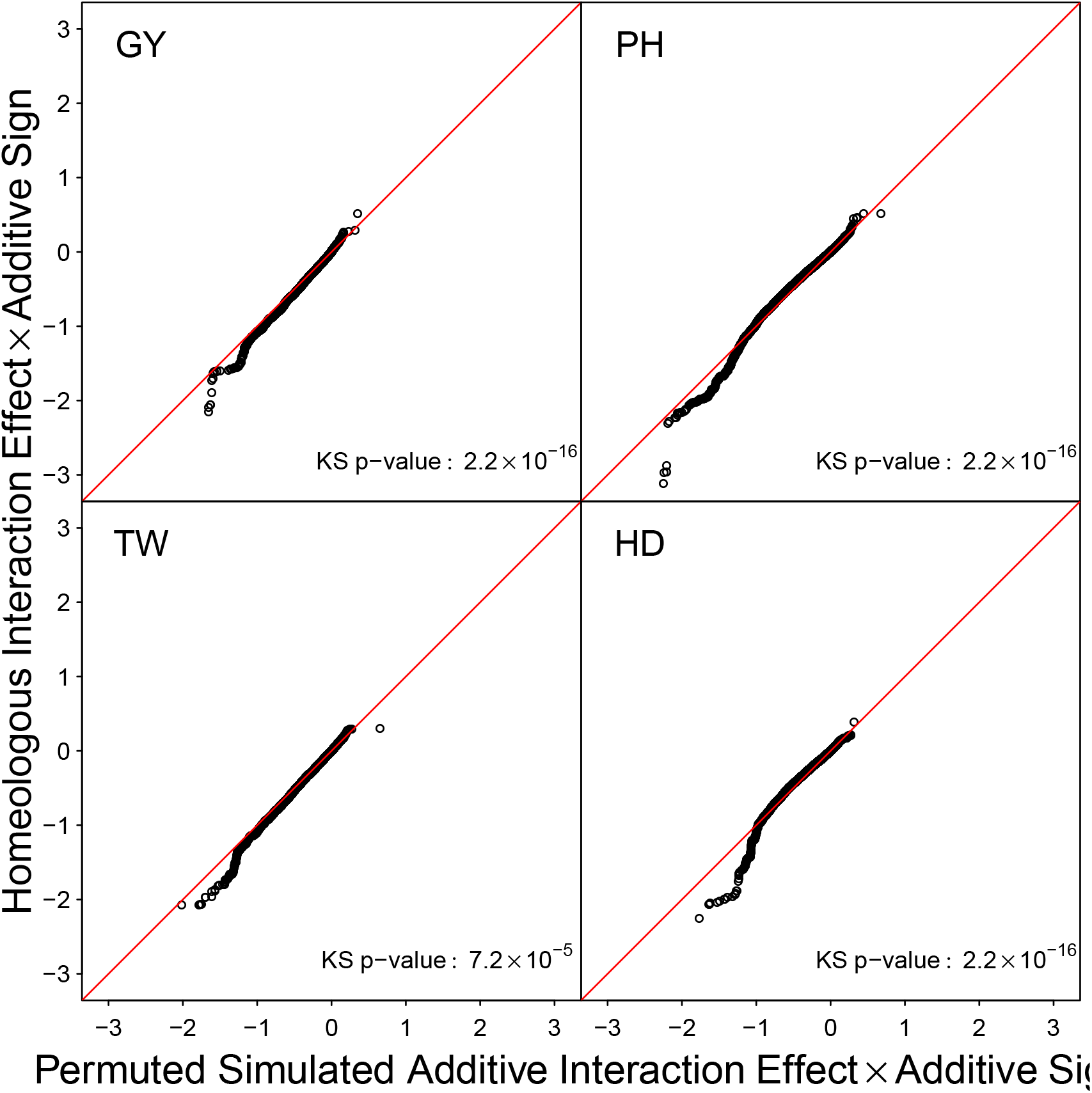
Quantile quantile plot of the ordered estimated homeologous interaction effects plotted against those from a simulated phenotype sampled to obtain no epistatic interactions using the HTEV marker orientation. Markers scores were permuted before simulation of the phenotype to remove LD between markers. Interaction effects have been multiplied by the effect sign of the corresponding additive effects to emphasize the relationship between the additive and interaction effects.

**Figure S7:**
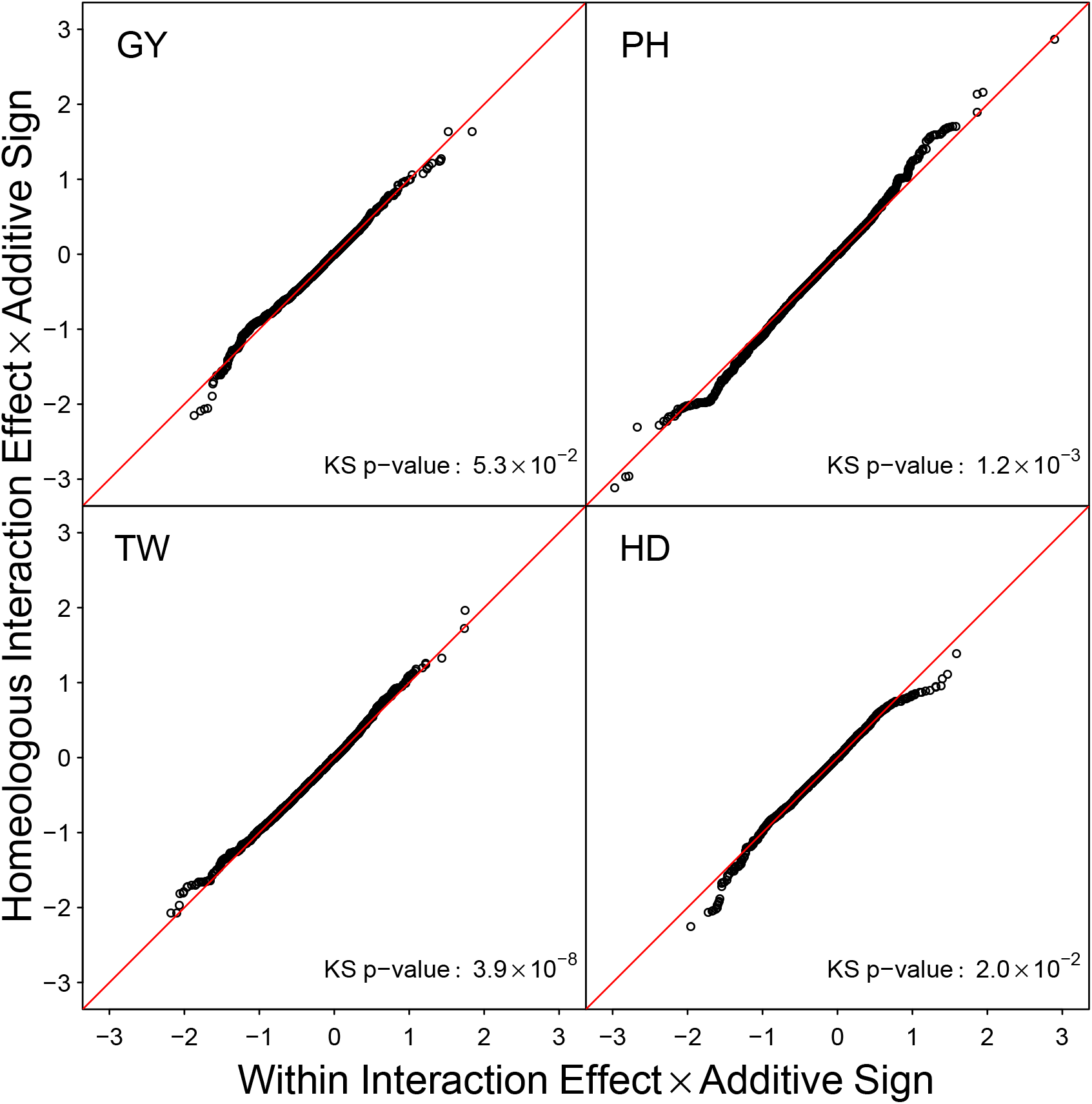
Quantile quantile plot of the ordered estimated homeologous interaction effects plotted against those from marker sets sampled within subgenome chromosomes (Within) using the LAVHAE. Interaction effects have been multiplied by the effect sign of the corresponding additive effects to emphasize the relationship between the additive and interaction effects.

**Figure S8:**
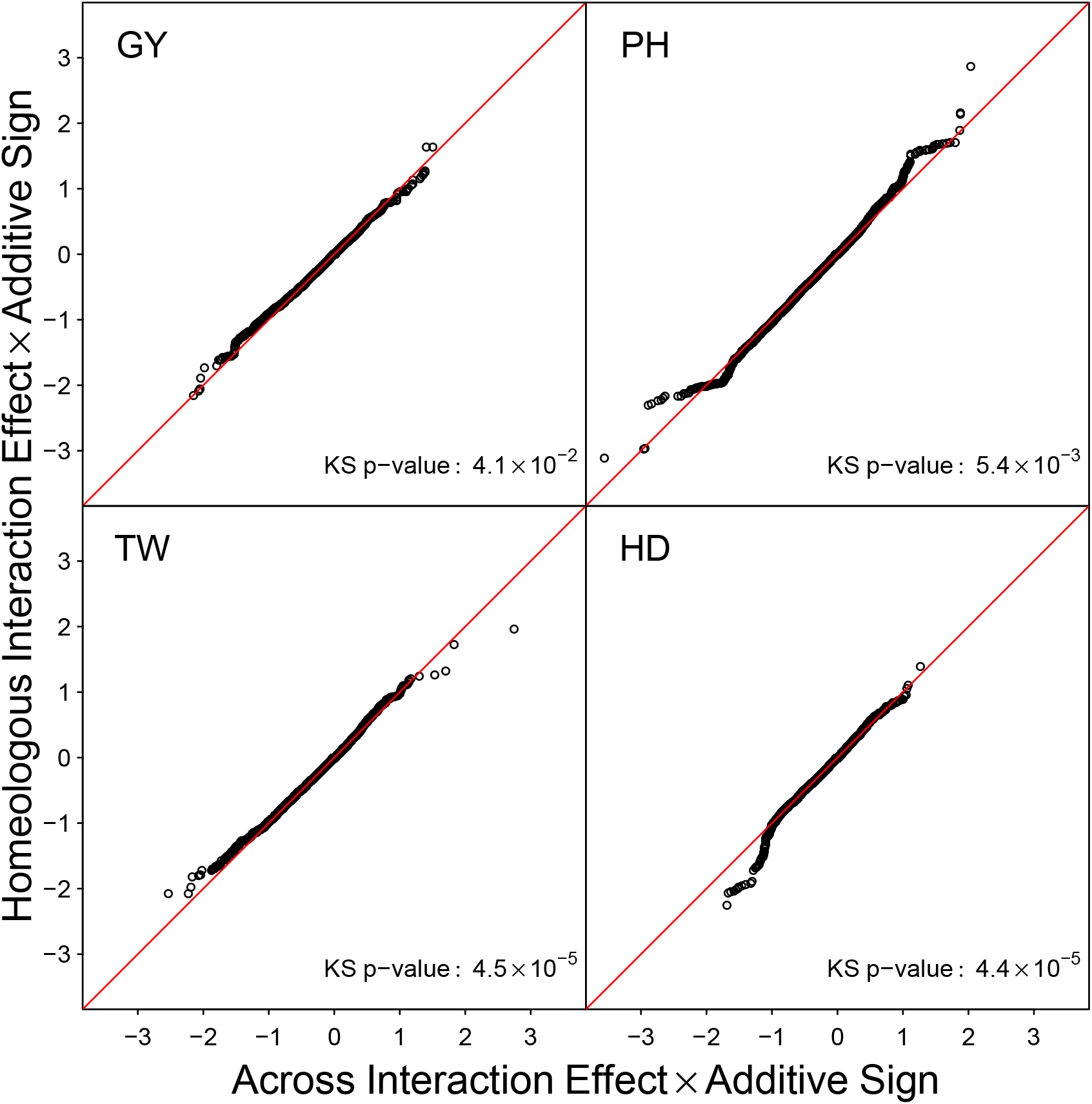
Quantile quantile plot of the ordered estimated homeologous interaction effects plotted against those from marker sets sampled across non-syntenic subgenome chromosomes (Across) using the LAVHAE marker orientation. Interaction effects have been multiplied by the effect sign of the corresponding additive effects to emphasize the relationship between the additive and interaction effects.

**Figure S9:**
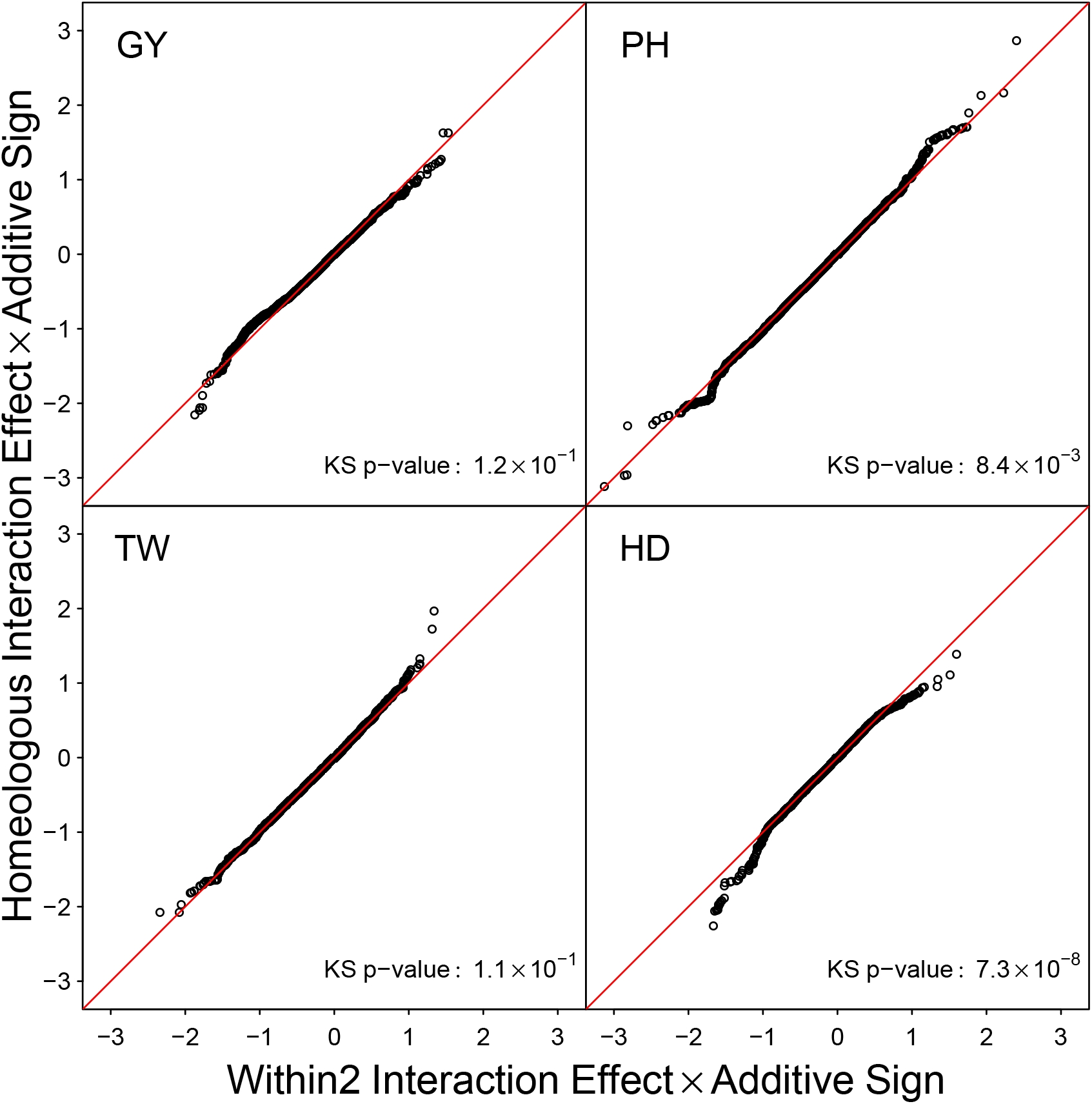
Quantile quantile plot of the ordered estimated homeologous interaction effects plotted against those from marker sets re-sampled within subgenome chromosomes (Within2) using the LAVHAE. Interaction effects have been multiplied by the effect sign of the corresponding additive effects to emphasize the relationship between the additive and interaction effects.

**Figure S10:**
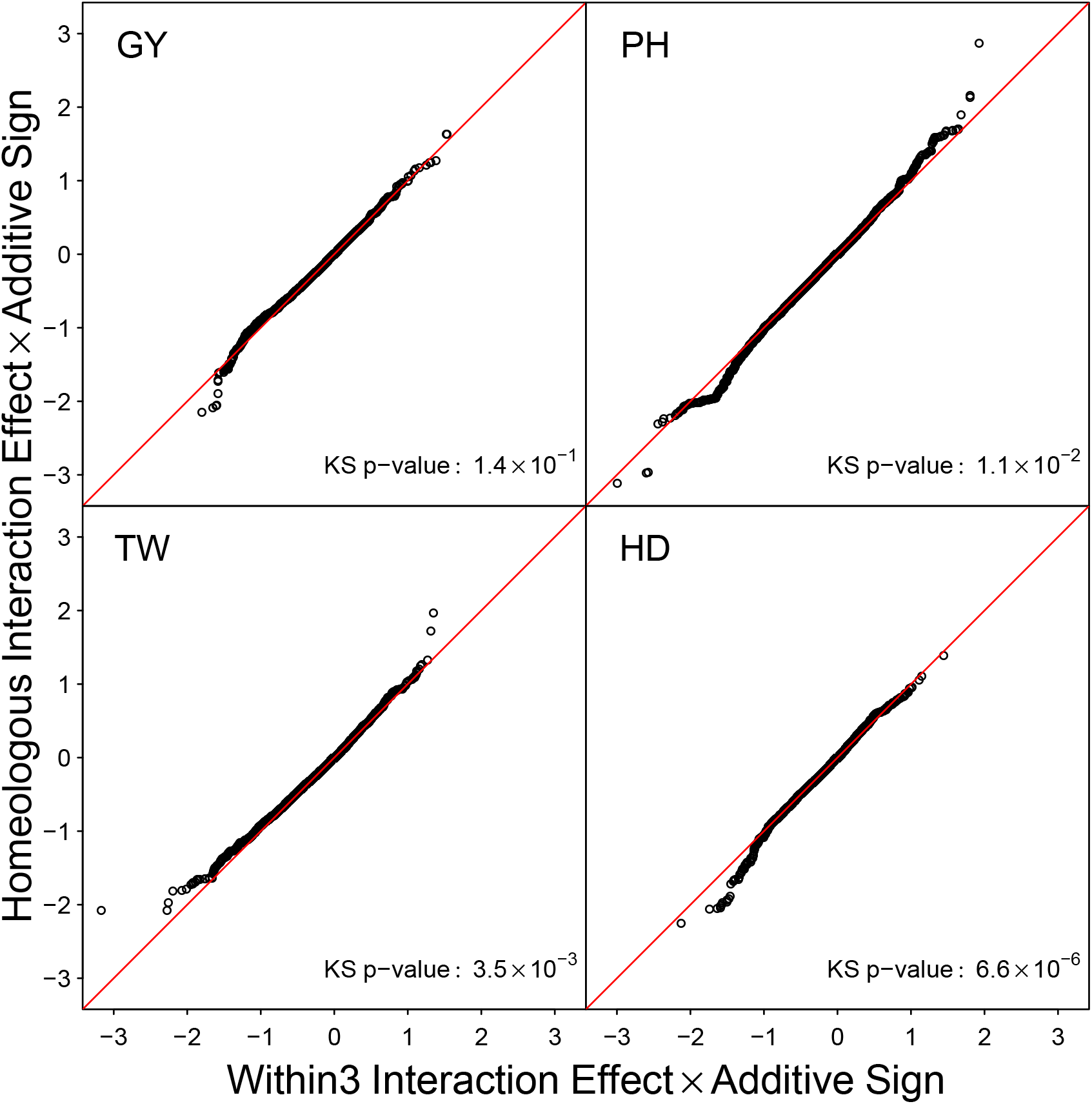
Quantile quantile plot of the ordered estimated homeologous interaction effects plotted against those from marker sets re-sampled within subgenome chromosomes (Within3) using the LAVHAE. Interaction effects have been multiplied by the effect sign of the corresponding additive effects to emphasize the relationship between the additive and interaction effects.

**Figure S11:**
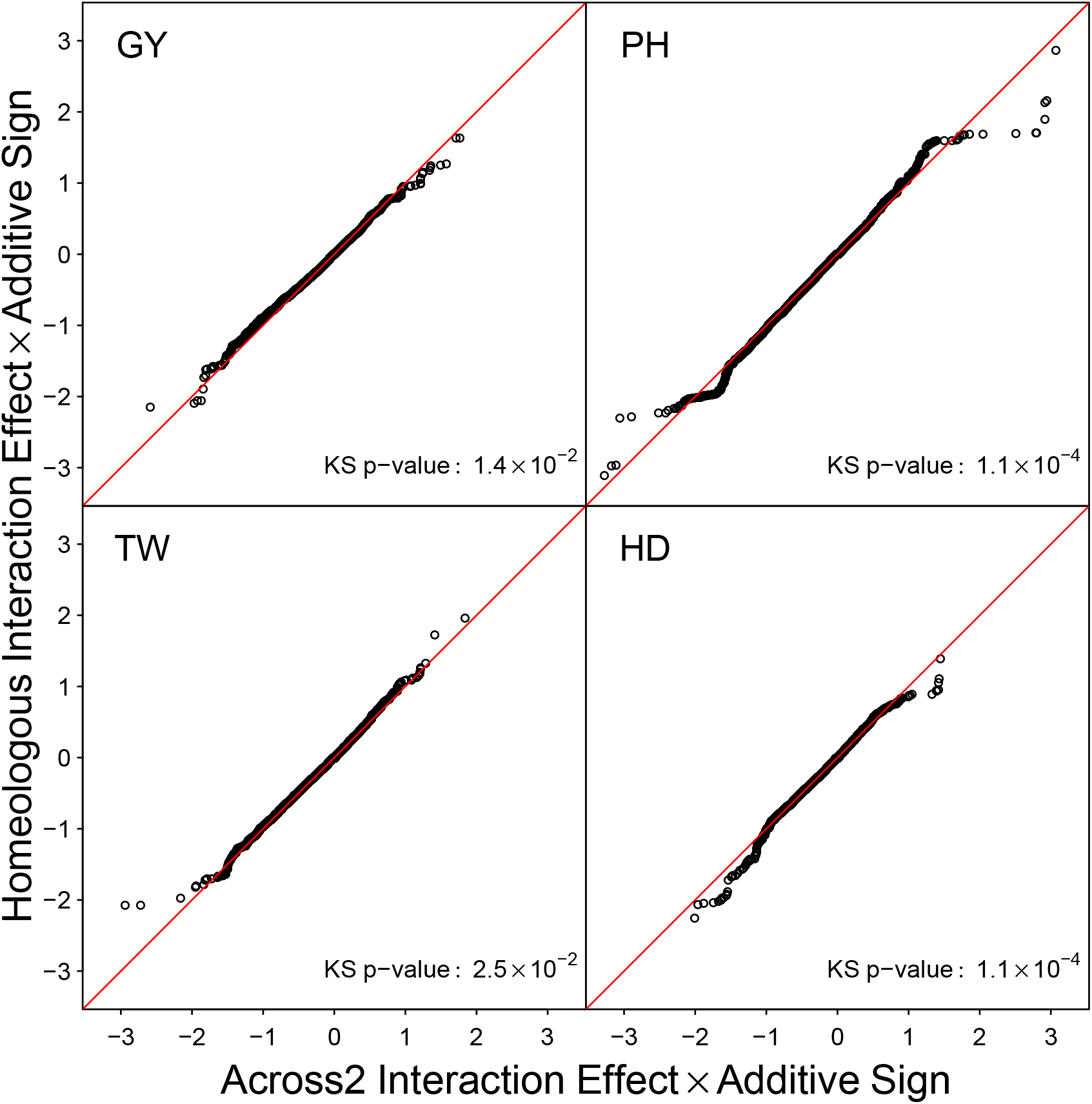
Quantile quantile plot of the ordered estimated homeologous interaction effects plotted against those from marker sets re-sampled across non-syntenic subgenome chromosomes (Across2) using the LAVHAE marker orientation. Interaction effects have been multiplied by the effect sign of the corresponding additive effects to emphasize the relationship between the additive and interaction effects.

**Figure S12:**
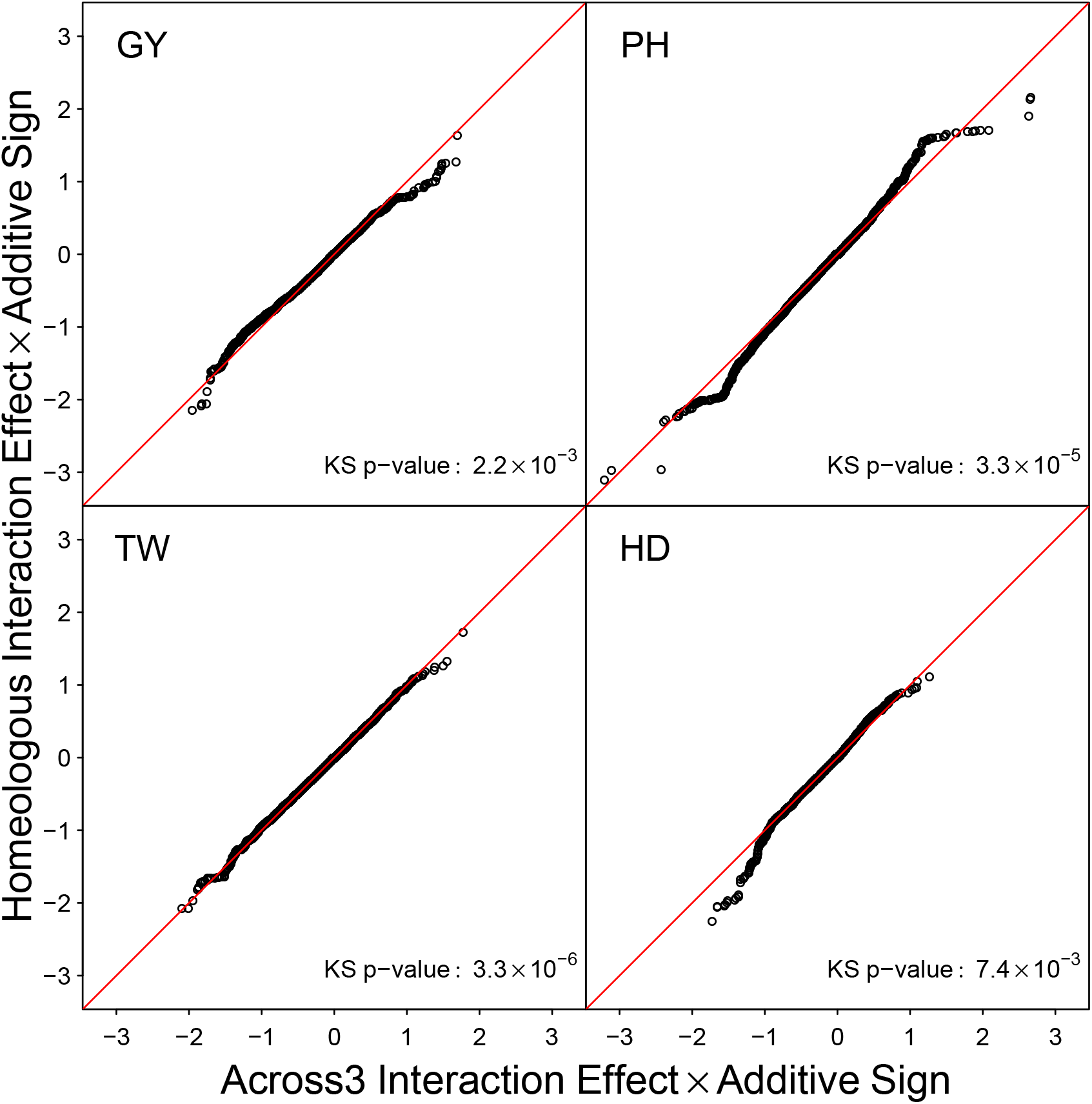
Quantile quantile plot of the ordered estimated homeologous interaction effects plotted against those from marker sets re-sampled across non-syntenic subgenome chromosomes (Across3) using the LAVHAE marker orientation. Interaction effects have been multiplied by the effect sign of the corresponding additive effects to emphasize the relationship between the additive and interaction effects.

**Table S4:**
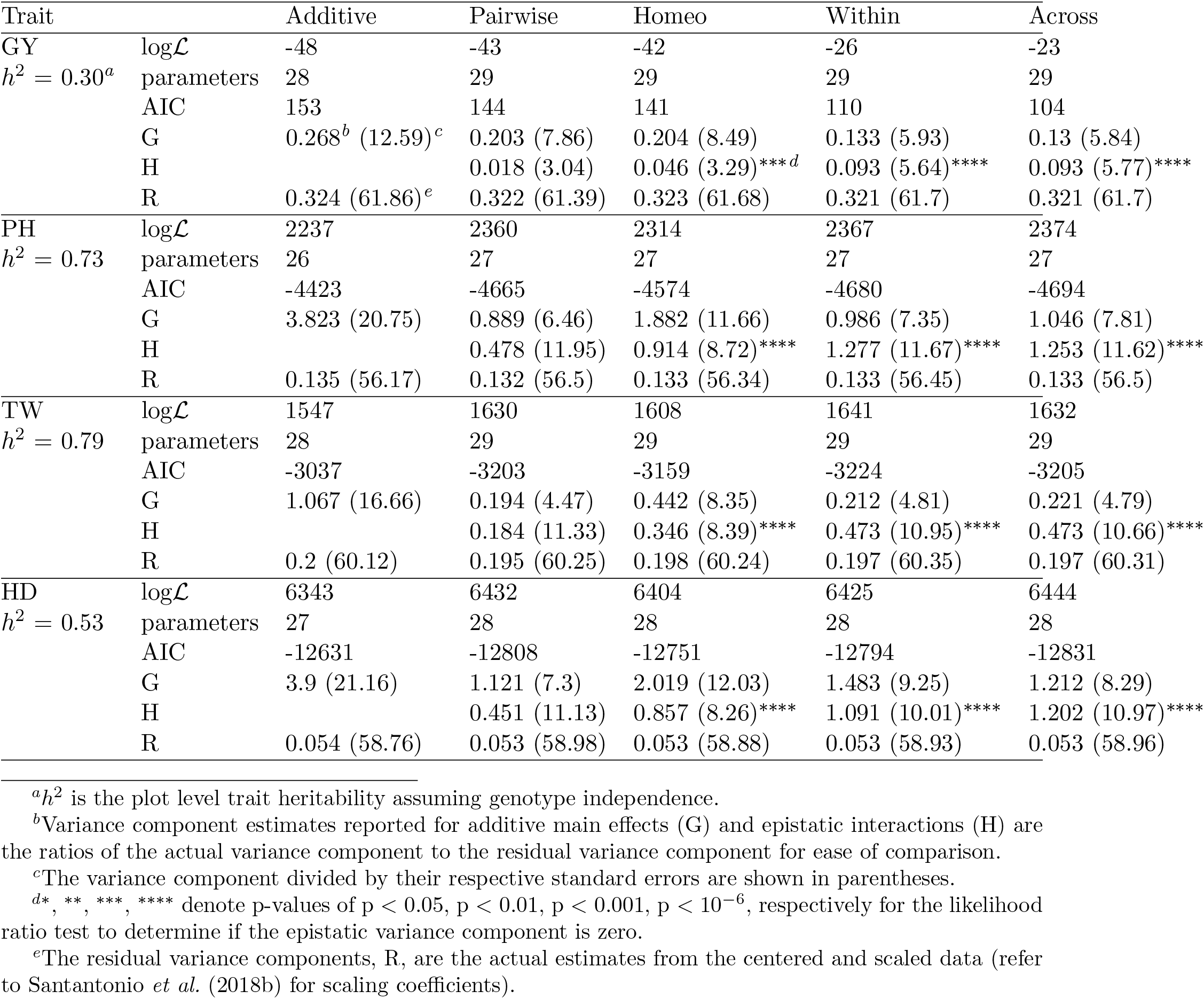
Mixed model REML fit summaries of one additive and four epistasis models for four traits (GY, PH, TW and HD) in the CNLM population based on the {−1, 1} marker parameterization using the LAVHAE marker orientation. Plot level heritabilities assuming genotype independence (i.i.d.) for each trait are shown underneath each trait name.

**Table S5:** Mixed model REML fit summaries of three epistasis models for 4 traits (GY, PH, TW and HD) in the CNLM population based on the {0, 1} marker parameterization using the LAVHAE marker orientation.

| Trait |  | Homeo | Within | Across |
| --- | --- | --- | --- | --- |
| GY | $\log\mathcal{L}$ | -48 | -47 | -42 |
|  | parameters | 29 | 29 | 29 |
|  | AIC | 155 | 152 | 143 |
|  | G | 0.267<sup>a</sup> (7.6)<sup>b</sup> | 0.207 (5.5) | 0.146 (4.16) |
|  | H | 0 (0.01) | 0.054 (1.73) | 0.108 (3.39)<sup>***c</sup> |
|  | R | 0.324 (61.81)<sup>d</sup> | 0.324 (61.77) | 0.324 (61.8) |
| PH | $\log\mathcal{L}$ | 2282 | 2268 | 2285 |
|  | parameters | 27 | 27 | 27 |
|  | AIC | -4510 | -4482 | -4516 |
|  | G | 1.198 (5.03) | 1.766 (6.95) | 1.177 (5.02) |
|  | H | 1.981 (8.36)<sup>****</sup> | 1.592 (6.95)<sup>****</sup> | 2.051 (8.66)<sup>****</sup> |
|  | R | 0.134 (56.23) | 0.134 (56.24) | 0.134 (56.24) |
| TW | $\log\mathcal{L}$ | 1560 | 1555 | 1567 |
|  | parameters | 29 | 29 | 29 |
|  | AIC | -3061 | -3052 | -3076 |
|  | G | 0.553 (5.88) | 0.659 (6.68) | 0.498 (5.57) |
|  | H | 0.414 (5.04)<sup>****</sup> | 0.331 (4.06)<sup>***</sup> | 0.482 (5.85)<sup>****</sup> |
|  | R | 0.199 (60.11) | 0.199 (60.1) | 0.198 (60.13) |
| HD | $\log\mathcal{L}$ | 6382 | 6364 | 6379 |
|  | parameters | 28 | 28 | 28 |
|  | AIC | -12709 | -12673 | -12702 |
|  | G | 1.51 (6.14) | 2.077 (7.82) | 1.659 (6.67) |
|  | H | 1.781 (7.73)<sup>****</sup> | 1.358 (6.09)<sup>****</sup> | 1.68 (7.36)<sup>****</sup> |
|  | R | 0.053 (58.84) | 0.054 (58.78) | 0.054 (58.81) |

<sup>a</sup>Variance component estimates reported for additive main effects (G) and epistatic interactions (H) are the ratios of the actual variance component to the residual variance component for ease of comparison.
<sup>b</sup>The variance component divided by their respective standard errors are shown in parentheses.
<sup>c</sup>\*, \*\*, \*\*\*, \*\*\*\* denote p-values of $p < 0.05$ , $p < 0.01$ , $p < 0.001$ , $p < 10^{-6}$ , respectively for the likelihood ratio test to determine if the epistatic variance component is zero.
<sup>d</sup>The residual variance components, R, are the actual estimates from the centered and scaled data (refer to Santantonio *et al.* (2018b) for scaling coefficients).

**Table S6:** Mixed model REML fit summaries of three epistasis models for 4 traits (GY, PH, TW and HD) in the CNLM population based on the {−1, 1} marker parameterization using the POS marker orientation.

| Trait |  | Homeo | Within | Across |
| --- | --- | --- | --- | --- |
| GY | $\log\mathcal{L}$ | -48 | -41 | -40 |
|  | parameters | 29 | 29 | 29 |
|  | AIC | 154 | 140 | 138 |
|  | G | 0.257<sup>a</sup> (10.31)<sup>b</sup> | 0.191 (7.44) | 0.186 (7.32) |
|  | H | 0.008 (0.75) | 0.052 (3.45)<sup>***c</sup> | 0.054 (3.61)<sup>***</sup> |
|  | R | 0.324 (61.7)<sup>d</sup> | 0.323 (61.64) | 0.323 (61.64) |
| PH | $\log\mathcal{L}$ | 2287 | 2323 | 2326 |
|  | parameters | 27 | 27 | 27 |
|  | AIC | -4521 | -4593 | -4598 |
|  | G | 2.316 (13.04) | 1.507 (9.34) | 1.551 (9.59) |
|  | H | 0.705 (7.3)<sup>****</sup> | 1.056 (9.85)<sup>****</sup> | 1.036 (9.72)<sup>****</sup> |
|  | R | 0.134 (56.29) | 0.133 (56.38) | 0.133 (56.4) |
| TW | $\log\mathcal{L}$ | 1589 | 1599 | 1604 |
|  | parameters | 29 | 29 | 29 |
|  | AIC | -3120 | -3139 | -3150 |
|  | G | 0.554 (9.49) | 0.437 (7.44) | 0.395 (7.02) |
|  | H | 0.282 (7.22)<sup>****</sup> | 0.354 (8.36)<sup>****</sup> | 0.368 (8.71)<sup>****</sup> |
|  | R | 0.198 (60.18) | 0.197 (60.2) | 0.197 (60.21) |
| HD | $\log\mathcal{L}$ | 6379 | 6393 | 6415 |
|  | parameters | 28 | 28 | 28 |
|  | AIC | -12701 | -12730 | -12774 |
|  | G | 2.547 (13.61) | 2.017 (10.81) | 1.689 (9.94) |
|  | H | 0.601 (6.43)<sup>****</sup> | 0.848 (8.04)<sup>****</sup> | 0.982 (9.26)<sup>****</sup> |
|  | R | 0.053 (58.83) | 0.053 (58.87) | 0.053 (58.92) |

<sup>a</sup>Variance component estimates reported for additive main effects (G) and epistatic interactions (H) are the ratios of the actual variance component to the residual variance component for ease of comparison.
<sup>b</sup>The variance component divided by their respective standard errors are shown in parentheses.
<sup>c\*</sup>, \*\*, \*\*\*, \*\*\*\* denote p-values of $p < 0.05$ , $p < 0.01$ , $p < 0.001$ , $p < 10^{-6}$ , respectively for the likelihood ratio test to determine if the epistatic variance component is zero.
<sup>d</sup>The residual variance components, R, are the actual estimates from the centered and scaled data (refer to Santantonio *et al.* (2018b) for scaling coefficients).

**Table S7:** Mixed model REML fit summaries of three epistasis models for 4 traits (GY, PH, TW and HD) in the CNLM population based on the {−1, 1} marker parameterization using the NEG marker orientation.

| Trait |  | Homeo | Within | Across |
| --- | --- | --- | --- | --- |
| GY | $\log\mathcal{L}$ | -46 | -38 | -35 |
|  | parameters | 29 | 29 | 29 |
|  | AIC | 151 | 134 | 129 |
|  | G | 0.236<sup>a</sup> (9.44)<sup>b</sup> | 0.181 (7.35) | 0.178 (7.35) |
|  | H | 0.022 (1.86) | 0.058 (3.9)<sup>***c</sup> | 0.06 (4.1)<sup>****</sup> |
|  | R | 0.324 (61.71)<sup>d</sup> | 0.323 (61.68) | 0.322 (61.68) |
| PH | $\log\mathcal{L}$ | 2293 | 2336 | 2342 |
|  | parameters | 27 | 27 | 27 |
|  | AIC | -4532 | -4619 | -4629 |
|  | G | 2.235 (12.79) | 1.428 (9.19) | 1.464 (9.46) |
|  | H | 0.746 (7.52)<sup>****</sup> | 1.061 (10.06)<sup>****</sup> | 1.038 (10.07)<sup>****</sup> |
|  | R | 0.134 (56.3) | 0.133 (56.39) | 0.133 (56.42) |
| TW | $\log\mathcal{L}$ | 1580 | 1605 | 1601 |
|  | parameters | 29 | 29 | 29 |
|  | AIC | -3101 | -3153 | -3144 |
|  | G | 0.614 (9.96) | 0.373 (6.78) | 0.388 (6.71) |
|  | H | 0.241 (6.4)<sup>****</sup> | 0.374 (8.94)<sup>****</sup> | 0.367 (8.59)<sup>****</sup> |
|  | R | 0.199 (60.15) | 0.198 (60.22) | 0.197 (60.21) |
| HD | $\log\mathcal{L}$ | 6380 | 6402 | 6409 |
|  | parameters | 28 | 28 | 28 |
|  | AIC | -12704 | -12747 | -12762 |
|  | G | 2.48 (13.41) | 1.88 (10.5) | 1.753 (10.09) |
|  | H | 0.626 (6.71)<sup>****</sup> | 0.895 (8.59)<sup>****</sup> | 0.95 (9.02)<sup>****</sup> |
|  | R | 0.053 (58.83) | 0.053 (58.89) | 0.053 (58.9) |

<sup>a</sup>Variance component estimates reported for additive main effects (G) and epistatic interactions (H) are the ratios of the actual variance component to the residual variance component for ease of comparison.
<sup>b</sup>The variance component divided by their respective standard errors are shown in parentheses.
<sup>c\*</sup>, <sup>\*\*</sup>, <sup>\*\*\*</sup>, <sup>\*\*\*\*</sup> denote p-values of $p < 0.05$ , $p < 0.01$ , $p < 0.001$ , $p < 10^{-6}$ , respectively for the likelihood ratio test to determine if the epistatic variance component is zero.
<sup>d</sup>The residual variance components, R, are the actual estimates from the centered and scaled data (refer to Santantonio *et al.* (2018b) for scaling coefficients).

**Table S8:** Mixed model REML fit summaries of three epistasis models for 4 traits (GY, PH, TW and HD) in the CNLM population based on the {−1, 1} marker parameterization using the HTEV marker orientation.

| trait |  | Homeo | Within | Across |
| --- | --- | --- | --- | --- |
| GY | $\log\mathcal{L}$ | -46 | -34 | -30 |
|  | parameters | 29 | 29 | 29 |
|  | AIC | 151 | 127 | 118 |
|  | G | 0.233<sup>a</sup> (9.23)<sup>b</sup> | 0.165 (6.86) | 0.151 (6.45) |
|  | H | 0.025 (1.97) | 0.071 (4.56)<sup>****c</sup> | 0.079 (5)<sup>****</sup> |
|  | R | 0.323 (61.65)<sup>d</sup> | 0.322 (61.66) | 0.322 (61.67) |
| PH | $\log\mathcal{L}$ | 2300 | 2355 | 2357 |
|  | parameters | 27 | 27 | 27 |
|  | AIC | -4546 | -4655 | -4659 |
|  | G | 2.052 (12.02) | 1.101 (7.81) | 1.142 (7.99) |
|  | H | 0.84 (8.12)<sup>****</sup> | 1.227 (11.24)<sup>****</sup> | 1.209 (11.09)<sup>****</sup> |
|  | R | 0.133 (56.32) | 0.133 (56.43) | 0.133 (56.46) |
| TW | $\log\mathcal{L}$ | 1599 | 1623 | 1623 |
|  | parameters | 29 | 29 | 29 |
|  | AIC | -3140 | -3189 | -3187 |
|  | G | 0.476 (8.51) | 0.283 (5.73) | 0.267 (5.4) |
|  | H | 0.335 (7.92)<sup>****</sup> | 0.435 (10.13)<sup>****</sup> | 0.45 (10.15)<sup>****</sup> |
|  | R | 0.198 (60.2) | 0.197 (60.29) | 0.197 (60.28) |
| HD | $\log\mathcal{L}$ | 6397 | 6410 | 6423 |
|  | parameters | 28 | 28 | 28 |
|  | AIC | -12738 | -12764 | -12790 |
|  | G | 2.13 (12.27) | 1.62 (9.54) | 1.395 (8.69) |
|  | H | 0.808 (7.9)<sup>****</sup> | 1.029 (9.43)<sup>****</sup> | 1.139 (10.18)<sup>****</sup> |
|  | R | 0.053 (58.88) | 0.053 (58.91) | 0.053 (58.94) |

<sup>a</sup>Variance component estimates reported for additive main effects (G) and epistatic interactions (H) are the ratios of the actual variance component to the residual variance component for ease of comparison.
<sup>b</sup>The variance component divided by their respective standard errors are shown in parentheses.
<sup>c</sup>\*, \*\*, \*\*\*, \*\*\*\* denote p-values of $p < 0.05$ , $p < 0.01$ , $p < 0.001$ , $p < 10^{-6}$ , respectively for the likelihood ratio test to determine if the epistatic variance component is zero.
<sup>d</sup>The residual variance components, R, are the actual estimates from the centered and scaled data (refer to Santantonio *et al.* (2018b) for scaling coefficients).

**Table S9:** Mixed model REML fit summaries of three epistasis models for 4 traits (GY, PH, TW and HD) in the CNLM population based on the {0, 1} marker parameterization using the HTEV marker orientation.

| trait |  | Homeo | Within | Across |
| --- | --- | --- | --- | --- |
| GY | $\log\mathcal{L}$ | -48 | -48 | -48 |
|  | parameters | 29 | 29 | 29 |
|  | AIC | 155 | 155 | 155 |
|  | G | 0.268<sup>a</sup> (12.59)<sup>b</sup> | 0.268 (12.59) | 0.268 (12.59) |
|  | H | 0 | 0 | 0 |
|  | R | 0.324 (61.86)<sup>c</sup> | 0.324 (61.86) | 0.324 (61.86) |
| PH | $\log\mathcal{L}$ | 2260 | 2246 | 2248 |
|  | parameters | 27 | 27 | 27 |
|  | AIC | -4466 | -4438 | -4443 |
|  | G | 1.981 (7.41) | 2.84 (9.98) | 2.502 (8.49) |
|  | H | 1.423 (6.05)<sup>****d</sup> | 0.806 (3.68)<sup>***</sup> | 1.081 (4.44)<sup>***</sup> |
|  | R | 0.134 (56.2) | 0.134 (56.19) | 0.134 (56.19) |
| TW | $\log\mathcal{L}$ | 1552 | 1547 | 1549 |
|  | parameters | 29 | 29 | 29 |
|  | AIC | -3046 | -3036 | -3041 |
|  | G | 0.746 (7.61) | 0.992 (9.51) | 0.857 (8.24) |
|  | H | 0.264 (3.38)<sup>***</sup> | 0.064 (0.87) | 0.183 (2.26)<sup>*</sup> |
|  | R | 0.199 (60.1) | 0.199 (60.08) | 0.199 (60.09) |
| HD | $\log\mathcal{L}$ | 6358 | 6350 | 6356 |
|  | parameters | 28 | 28 | 28 |
|  | AIC | -12660 | -12643 | -12656 |
|  | G | 2.528 (9.24) | 2.937 (10.1) | 2.468 (8.5) |
|  | H | 1.052 (4.83)<sup>****</sup> | 0.749 (3.44)<sup>***</sup> | 1.16 (4.78)<sup>****</sup> |
|  | R | 0.054 (58.79) | 0.054 (58.76) | 0.054 (58.78) |

<sup>a</sup>Variance component estimates reported for additive main effects (G) and epistatic interactions (H) are the ratios of the actual variance component to the residual variance component for ease of comparison.
<sup>b</sup>The variance component divided by their respective standard errors are shown in parentheses.
<sup>c</sup>The residual variance components, R, are the actual estimates from the centered and scaled data (refer to Santantonio *et al.* (2018b) for scaling coefficients).
<sup>d</sup>\*, \*\*, \*\*\*, \*\*\*\* denote p-values of $p < 0.05$ , $p < 0.01$ , $p < 0.001$ , $p < 10^{-6}$ , respectively for the likelihood ratio test to determine if the epistatic variance component is zero.

**Table S10:** Mixed model REML fit summaries of four epistasis models for 4 traits (GY, PH, TW and HD) in the CNLM population based on the {−1, 1} marker parameterization using the LAVHAE marker orientation using two additional samples of Within (Within2, Within3) and Across (Across2, Across3).

| Trait |  | Within2 | Within3 | Across2 | Across3 |
| --- | --- | --- | --- | --- | --- |
| GY | $\log\mathcal{L}$ | -32 | -29 | -26 | -32 |
|  | parameters | 29 | 29 | 29 | 29 |
|  | AIC | 122 | 115 | 110 | 121 |
|  | G | 0.15<sup>a</sup> (6.29)<sup>b</sup> | 0.14 (6.07) | 0.134 (5.91) | 0.154 (6.55) |
|  | H | 0.081 (4.97)<sup>****c</sup> | 0.086 (5.32)<sup>****</sup> | 0.092 (5.58)<sup>****</sup> | 0.078 (4.89)<sup>****</sup> |
|  | R | 0.322 (61.67)<sup>d</sup> | 0.322 (61.7) | 0.321 (61.69) | 0.322 (61.67) |
| PH | $\log\mathcal{L}$ | 2369 | 2356 | 2357 | 2359 |
|  | parameters | 27 | 27 | 27 | 27 |
|  | AIC | -4685 | -4658 | -4661 | -4664 |
|  | G | 0.979 (7.38) | 1.066 (7.57) | 1.083 (7.67) | 1.103 (7.86) |
|  | H | 1.257 (11.59)<sup>****</sup> | 1.242 (11.23)<sup>****</sup> | 1.253 (11.27)<sup>****</sup> | 1.23 (11.27)<sup>****</sup> |
|  | R | 0.133 (56.45) | 0.133 (56.44) | 0.133 (56.44) | 0.133 (56.48) |
| TW | $\log\mathcal{L}$ | 1647 | 1627 | 1641 | 1634 |
|  | parameters | 29 | 29 | 29 | 29 |
|  | AIC | -3235 | -3196 | -3224 | -3210 |
|  | G | 0.2 (4.56) | 0.267 (5.42) | 0.213 (4.74) | 0.233 (5) |
|  | H | 0.479 (10.94)<sup>****</sup> | 0.441 (10.11)<sup>****</sup> | 0.48 (10.86)<sup>****</sup> | 0.472 (10.67)<sup>****</sup> |
|  | R | 0.197 (60.38) | 0.197 (60.29) | 0.196 (60.31) | 0.196 (60.36) |
| HD | $\log\mathcal{L}$ | 6455 | 6429 | 6442 | 6444 |
|  | parameters | 28 | 28 | 28 | 28 |
|  | AIC | -12853 | -12801 | -12828 | -12832 |
|  | G | 1.053 (7.75) | 1.366 (8.76) | 1.174 (8.15) | 1.173 (8.15) |
|  | H | 1.311 (11.68)<sup>****</sup> | 1.15 (10.39)<sup>****</sup> | 1.248 (11.19)<sup>****</sup> | 1.226 (11.11)<sup>****</sup> |
|  | R | 0.053 (58.99) | 0.053 (58.94) | 0.053 (58.94) | 0.053 (58.95) |

<sup>a</sup>Variance component estimates reported for additive main effects (G) and epistatic interactions (H) are the ratios of the actual variance component to the residual variance component for ease of comparison.
<sup>b</sup>The variance component divided by their respective standard errors are shown in parentheses.
<sup>c</sup>\*, \*\*, \*\*\*, \*\*\*\* denote p-values of $p < 0.05$ , $p < 0.01$ , $p < 0.001$ , $p < 10^{-6}$ , respectively for the likelihood ratio test to determine if the epistatic variance component is zero.
<sup>d</sup>The residual variance components, R, are the actual estimates from the centered and scaled data (refer to Santantonio *et al.* (2018b) for scaling coefficients).

**Table S11:**
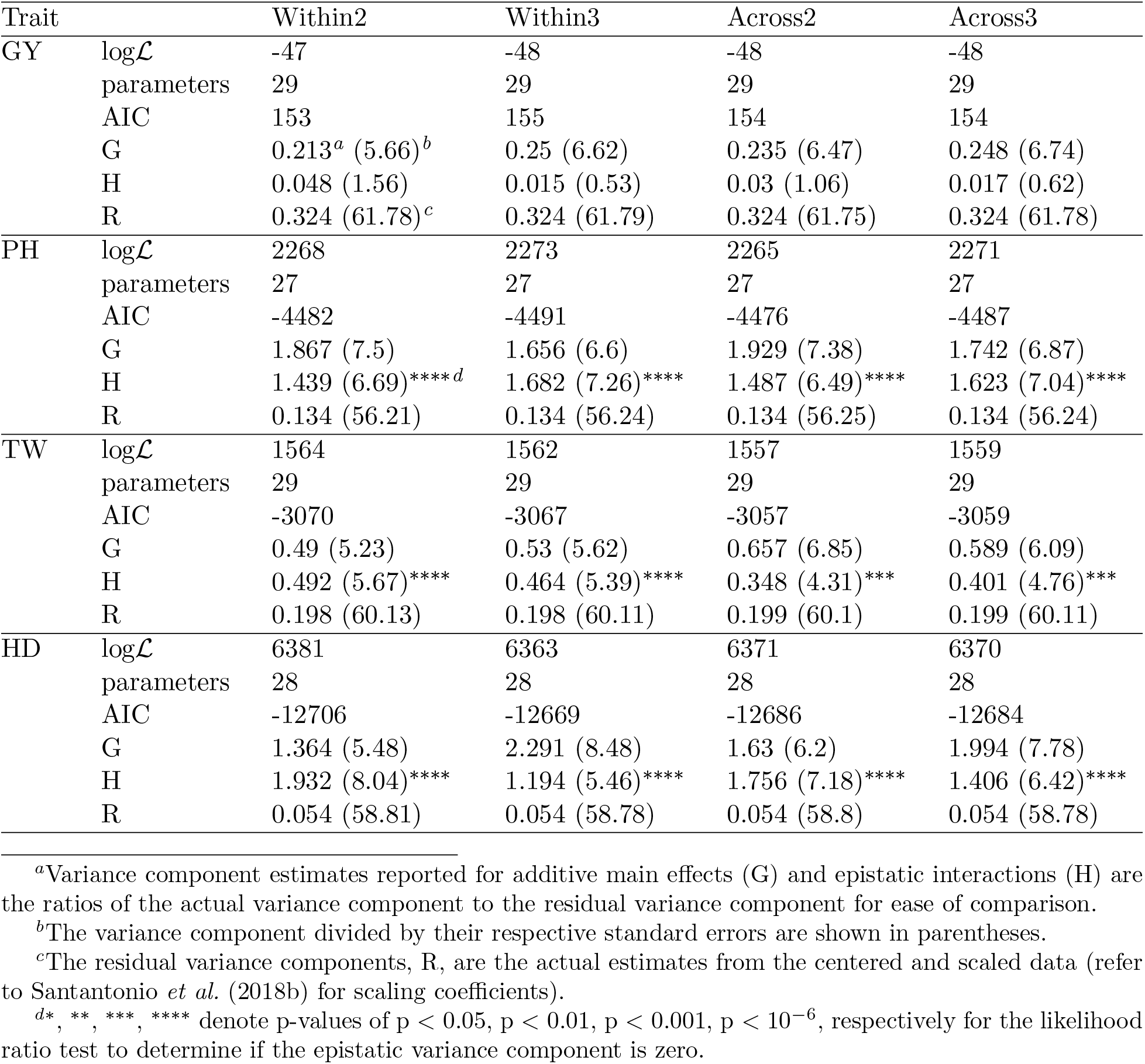
Mixed model REML fit summaries of four epistasis models for 4 traits (GY, PH, TW and HD) in the CNLM population based on the {0, 1} marker parameterization using the LAVHAE marker orientation using two additional samples of within and across markers.

**Table S12:**
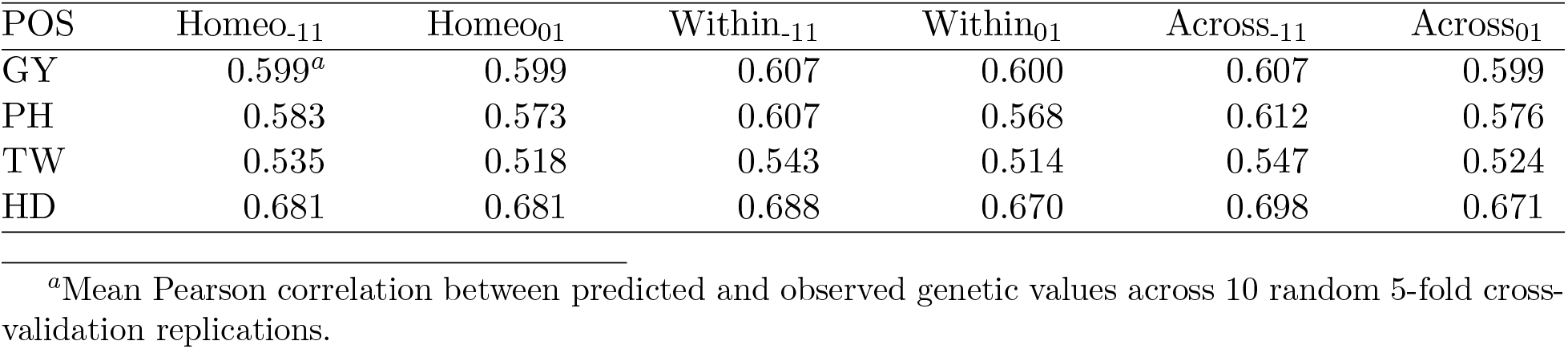
Prediction accuracies of Homeo, Within and Across genome marker sets for both {−1, 1} and {0, 1} marker coding using POS marker orientation.

| POS | Homeo<sub>-11</sub> | Homeo<sub>01</sub> | Within<sub>-11</sub> | Within<sub>01</sub> | Across<sub>-11</sub> | Across<sub>01</sub> |
| --- | --- | --- | --- | --- | --- | --- |
| GY | 0.599<sup>a</sup> | 0.599 | 0.607 | 0.600 | 0.607 | 0.599 |
| PH | 0.583 | 0.573 | 0.607 | 0.568 | 0.612 | 0.576 |
| TW | 0.535 | 0.518 | 0.543 | 0.514 | 0.547 | 0.524 |
| HD | 0.681 | 0.681 | 0.688 | 0.670 | 0.698 | 0.671 |

<sup>a</sup>Mean Pearson correlation between predicted and observed genetic values across 10 random 5-fold cross-validation replications.

**Table S13:** Prediction accuracies of Homeo, Within and Across genome marker sets for both {−1, 1} and {0, 1} marker coding using NEG marker orientation.

| NEG | Homeo<sub>-11</sub> | Homeo<sub>01</sub> | Within<sub>-11</sub> | Within<sub>01</sub> | Across<sub>-11</sub> | Across<sub>01</sub> |
| --- | --- | --- | --- | --- | --- | --- |
| GY | 0.602<sup>a</sup> | 0.599 | 0.612 | 0.599 | 0.615 | 0.600 |
| PH | 0.589 | 0.582 | 0.620 | 0.565 | 0.615 | 0.579 |
| TW | 0.535 | 0.513 | 0.555 | 0.510 | 0.546 | 0.519 |
| HD | 0.676 | 0.671 | 0.698 | 0.671 | 0.697 | 0.680 |

<sup>a</sup>Mean Pearson correlation between predicted and observed genetic values across 10 random 5-fold cross-validation replications.

**Table S14:** Prediction accuracies of Homeo, Within and Across genome marker sets for both {−1, 1} and {0, 1} marker coding using HTEV marker orientation.

| HTEV | Homeo<sub>-11</sub> | Homeo<sub>01</sub> | Within<sub>-11</sub> | Within<sub>01</sub> | Across<sub>-11</sub> | Across<sub>01</sub> |
| --- | --- | --- | --- | --- | --- | --- |
| GY | 0.601<sup>a</sup> | 0.601 | 0.616 | 0.600 | 0.621 | 0.600 |
| PH | 0.591 | 0.565 | 0.640 | 0.557 | 0.633 | 0.558 |
| TW | 0.548 | 0.513 | 0.572 | 0.513 | 0.568 | 0.513 |
| HD | 0.688 | 0.669 | 0.700 | 0.666 | 0.706 | 0.667 |

<sup>a</sup>Mean Pearson correlation between predicted and observed genetic values across 10 random 5-fold cross-validation replications.

**Table S15:** Prediction accuracies of two additional samples of Within (Within2, Within3) and Across (Across2, Across3) genome marker sets, for both {−1, 1} and {0, 1} marker coding using LAVHAE marker orientation.

| LAVHAE | Within2<sub>01</sub> | Within3<sub>01</sub> | Across2<sub>01</sub> | Across3<sub>01</sub> | Within2<sub>-11</sub> | Within3<sub>-11</sub> | Across2<sub>-11</sub> | Across3<sub>-11</sub> |
| --- | --- | --- | --- | --- | --- | --- | --- | --- |
| GY | 0.600 | 0.599 | 0.600 | 0.600 | 0.620 | 0.624 | 0.623 | 0.618 |
| PH | 0.573 | 0.569 | 0.566 | 0.570 | 0.655 | 0.640 | 0.634 | 0.644 |
| TW | 0.522 | 0.524 | 0.518 | 0.518 | 0.604 | 0.581 | 0.592 | 0.585 |
| HD | 0.683 | 0.673 | 0.676 | 0.679 | 0.727 | 0.715 | 0.718 | 0.724 |

**Table S16:** Estimates of *s* coefficients for marker sets where both additive and the two-way interaction effects were significant at *p* < 0.05 for each of 4 traits. The expected number of non-zero additive and two-way interactions effects based on a 0.05 significance threshold by chance for each trait is 3 (i.e. 22, 411 two-way interactions × 0.05^3^). Coefficients have been grouped by categories related to the potential mode of epistasis, where *s* < 0.5 indicates a highly negative interaction, 0.5 ≤ *s* < 1 a less-than-additive interaction may be indicative of subfunctionalization for homeologous genes, and *s* > 1 which indicates positive, or greater-than-additive, epistasis. Three marker sets are shown, either across all homeologous loci (Homeo), sampled sets within (Within) and across (Across) non-syntenic subgenome regions. An additional phenotype was simulated to contain additive only phenotypes to contain no epistasis, and fit with the Homeo marker set (Simulated Additive).

| Marker Set | Trait | $s < 0.5$ | $0.5 \leq s < 1$ | $s > 1$ | Total<sup>a</sup> |
| --- | --- | --- | --- | --- | --- |
| Homeo | GY | 0 | 0 | 2 | 2 |
| Homeo | PH | 2 | 8 | 4 | 14*** |
| Homeo | TW | 5 | 4 | 2 | 11** |
| Homeo | HD | 1 | 2 | 0 | 3 |
| Simulated Additive | PH | 0 | 1 | 3 | 4 |
| Simulated Additive | HD | 1 | 0 | 1 | 2 |
| Across | GY | 2 | 2 | 1 | 5 |
| Across | PH | 3 | 0 | 0 | 3 |
| Across | TW | 2 | 1 | 0 | 3 |
| Across | HD | 2 | 4 | 0 | 6* |
| Within | GY | 1 | 1 | 0 | 2 |
| Within | PH | 2 | 1 | 3 | 6* |
| Within | TW | 1 | 1 | 1 | 3 |
| Within | HD | 2 | 0 | 0 | 2 |

<sup>a</sup>\*, \*\*, \*\*\* indicate significantly greater than the expected number of significant sets at $p = 0.05$ , $10^{-4}$ and $10^{-6}$ based the binomial distribution with 22,411 trials and a probability of $0.05^3$ .

**Figure S13:**
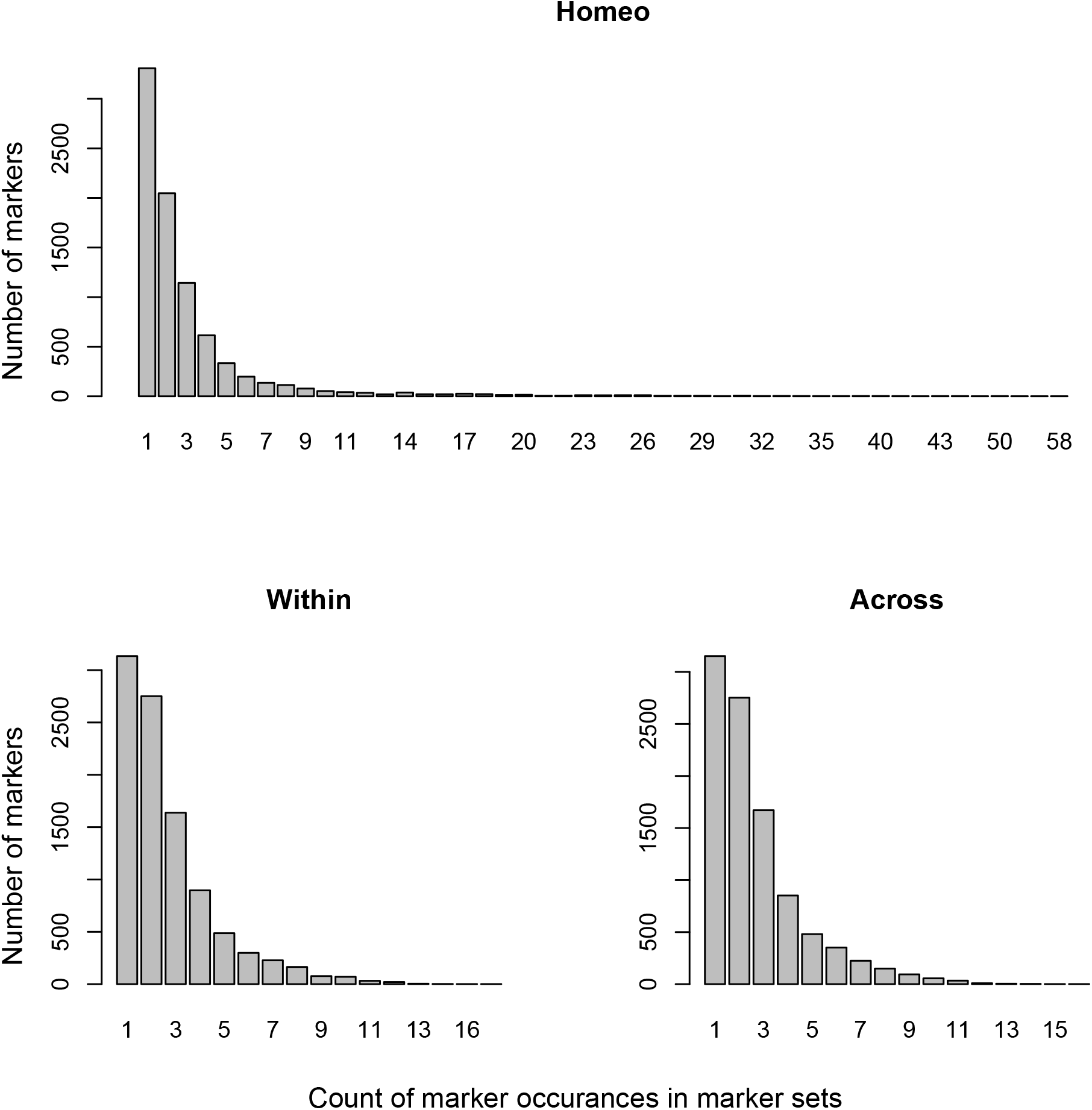
Distribution of the number of marker occurrences in marker sets. An occurrence of 1 indicates that a marker was only included in one marker set, whereas an occurrence of 10 would indicate that the marker was included in 10 marker sets.

**Figure S14:**
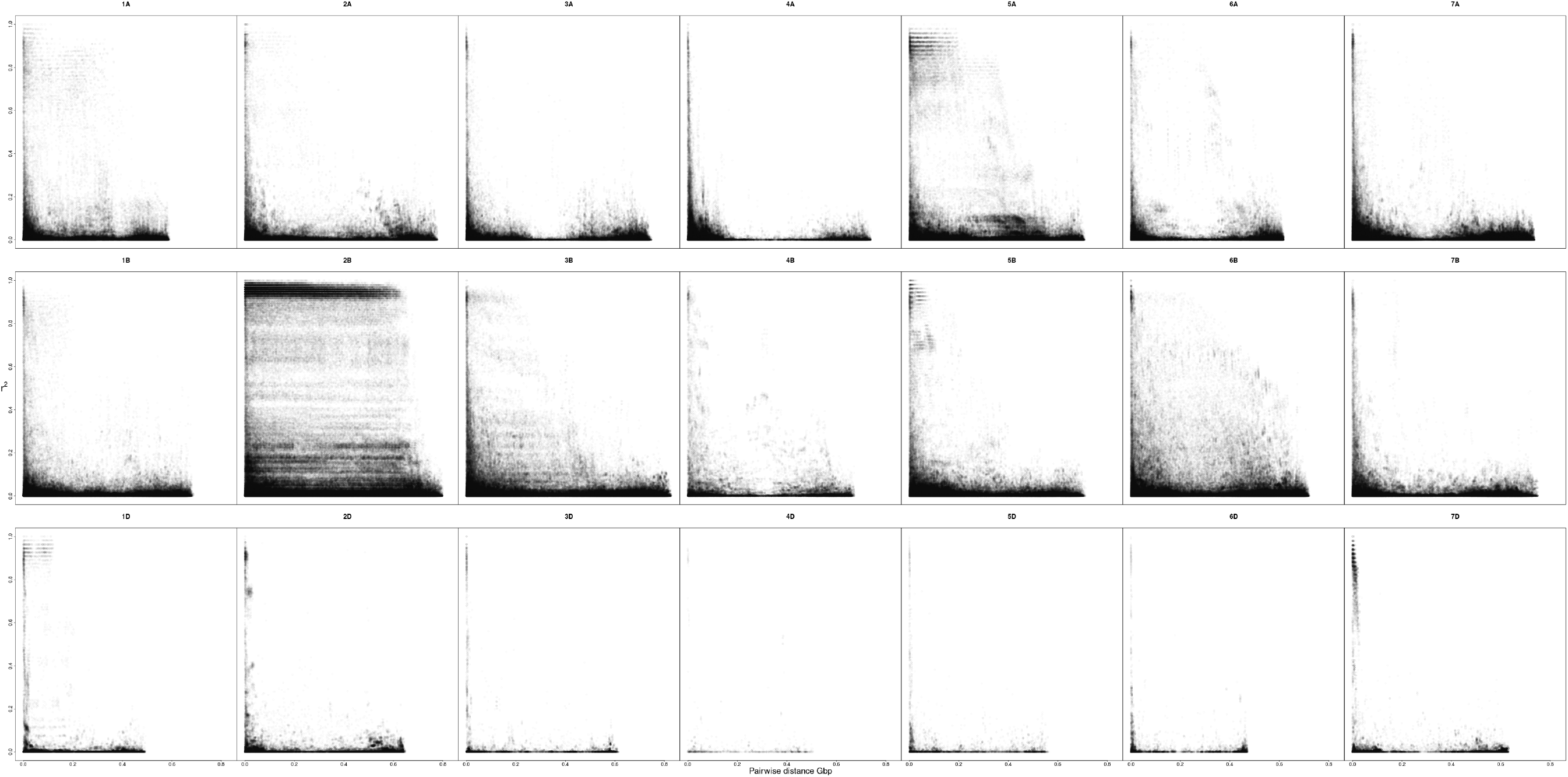
Pairwise linkage disequilibrium *r*^2^ values for the 21 wheat chromosomes in the CNLM population.

